# Insights from the IronTract challenge: optimal methods for mapping brain pathways from multi-shell diffusion MRI

**DOI:** 10.1101/2021.12.17.472836

**Authors:** Chiara Maffei, Gabriel Girard, Kurt G. Schilling, Dogu Baran Aydogan, Nagesh Adluru, Andrey Zhylka, Ye Wu, Matteo Mancini, Andac Hamamci, Alessia Sarica, Achille Teillac, Steven H. Baete, Davood Karimi, Fang-Cheng Yeh, Mert E. Yildiz, Ali Gholipour, Yann Bihan-Poudec, Bassem Hiba, Andrea Quattrone, Aldo Quattrone, Tommy Boshkovski, Nikola Stikov, Pew-Thian Yap, Alberto de Luca, Josien Pluim, Alexander Leemans, Vivek Prabhakaran, Barbara B. Bendlin, Andrew L. Alexander, Bennett A. Landman, Erick J. Canales-Rodríguez, Muhamed Barakovic, Jonathan Rafael-Patino, Thomas Yu, Gaëtan Rensonnet, Simona Schiavi, Alessandro Daducci, Marco Pizzolato, Elda Fischi-Gomez, Jean-Philippe Thiran, George Dai, Giorgia Grisot, Nikola Lazovski, Santi Puch, Marc Ramos, Paulo Rodrigues, Vesna Prchkovska, Robert Jones, Julia Lehman, Suzanne N. Haber, Anastasia Yendiki

## Abstract

Limitations in the accuracy of brain pathways reconstructed by diffusion MRI (dMRI) tractography have received considerable attention. While the technical advances spearheaded by the Human Connectome Project (HCP) led to significant improvements in dMRI data quality, it remains unclear how these data should be analyzed to maximize tractography accuracy. Over a period of two years, we have engaged the dMRI community in the IronTract Challenge, which aims to answer this question by leveraging a unique dataset. Macaque brains that have received both tracer injections and *ex vivo* dMRI at high spatial and angular resolution allow a comprehensive, quantitative assessment of tractography accuracy on state-of-the-art dMRI acquisition schemes. We find that, when analysis methods are carefully optimized, the HCP scheme can achieve similar accuracy as a more time-consuming, Cartesian-grid scheme. Importantly, we show that simple pre- and post-processing strategies can improve the accuracy and robustness of many tractography methods. Finally, we find that fiber configurations that go beyond crossing (*e*.*g*., fanning, branching) are the most challenging for tractography. The IronTract Challenge remains open and we hope that it can serve as a valuable validation tool for both users and developers of dMRI analysis methods.

## Introduction

Diffusion MRI (dMRI) tractography allows us to image brain pathways *in vivo* and non-invasively, and is thus a useful tool in a variety of research and clinical settings. However, it relies on indirect measurements of axonal orientations extracted from the dMRI signal, which can lead to errors in the reconstructed pathways. Possible sources of these errors, as identified by early studies, included uncertainty in the signal due to imaging noise^1^ and crossing fibers^2^. These issues motivated the effort to improve the signal-to-noise ratio (SNR), as well as the spatial and angular resolution of dMRI. The Human Connectome Project (HCP) sought to address these needs by developing scanners with ultra-high gradients, which allowed higher b-values to be acquired without sacrificing SNR, and accelerated dMRI sequences, which enabled higher angular and spatial resolution with shorter acquisition times^3,4^. These developments made multi-shell dMRI data prevalent. In parallel, orientation reconstruction methods were adapted to make better use of such data^5–9^.

These advances in data acquisition and analysis improved our ability to resolve crossing fibers within a voxel^10,11^ and allowed us to reconstruct white-matter circuitry in greater detail than previously possible^12,13^. However, it is unclear which analysis methods maximize the anatomic accuracy of the pathways that can be reconstructed from these state-of-the-art acquisition protocols. Given the large amounts of HCP-style, multi-shell data that are now publicly available^3,14–16^, and the plethora of methods for pre-processing, orientation reconstruction, and tractography that can be applied to these data, it is of critical importance to compare these methods with respect to objective metrics of anatomic accuracy.

Anatomic tracing in non-human primates (NHPs) can be used to assess the accuracy of tractography in the brain^17^. It allows us to reconstruct the complete trajectories of axon bundles from a tracer injection site to their destinations throughout the brain. The majority of previous studies that compared dMRI tractography to anatomic tracing were limited to single-shell dMRI data^18–24^. Furthermore, the majority of such studies only considered the end points of the fiber bundles, and not their complete trajectory^23–28^. That is because they did not have dMRI and tracer data from the same brains, hence they relied on connectivity matrices from existing tracer databases.

The IronTract Challenge is the first open tractography challenge to be conducted on high-resolution, densely sampled brain dMRI data. This allowed us to evaluate tractography accuracy for two widely adopted sampling schemes: multi-shell and Cartesian-grid. We leveraged a unique collection of NHP brains, where both anatomic tracer injections and *ex vivo* dMRI had been performed^29–31^. The availability of dMRI and tracer data in the same brains allowed us to evaluate the accuracy of tractography not only at the end points of the axon bundles but along their trajectory in the white matter. This is the only way localize exactly *where* tractography algorithms go wrong, which is a necessary step towards determining *why* they go wrong, and therefore how to improve them.

The IronTract Challenge also differed from previous tractography challenges in terms of its design. Participants submitted results with a wide range of tractography thresholds. When methods are compared only at their default thresholds (*e*.*g*., ^19,22,32^), they differ in terms of both sensitivity and specificity, and it is impossible to disentangle the effect of the threshold and the effect of the algorithm. Our design allowed us to circumvent this issue and to compare algorithms in terms of their sensitivity at the same level of specificity.

A previous validation study used data only from the training case of this challenge and performed a systematic comparison of a small number of q-space sampling, orientation reconstruction, and tractography methods, in all their permutations^29^. The IronTract Challenge expands the scope of our prior validation studies in two major ways. First, challenge participants chose a much wider range of state-of-the-art orientation reconstruction and tractography methods. Second, the addition of the validation case, which involved an injection in a different anatomical location and fibers following very different trajectories than the training case, allowed us to compare the robustness of the methods to the location of the seed region.

The IronTract Challenge was administered in two rounds (https://irontract.mgh.harvard.edu). The first round was organized in the context of the 2019 international conference on Medical Image Computing and Computer-Assisted Intervention. Preliminary results from the first and second rounds were presented, respectively, at the 2020 and 2021 annual meetings of the International Society for Magnetic Resonance in Medicine^33,34^. In the first round, two teams outperformed all others, achieving both high accuracy and robustness to the location of the seed region. This motivated the second round, where all participants revisited their analyses, replacing their pre- and post-processing steps with those of the two high-performing teams. This allowed us to investigate the extent to which performance was dependent on the pre- and post-processing vs. the orientation reconstruction and tractography methods. The outcomes of this effort, as detailed below, include *(i)* practical recommendations for users of HCP-style, multi-shell dMRI data, who are interested in methods for analyzing these data that maximize anatomical accuracy, and *(ii)* insights on the fundamental failure modes of tractography for method developers, who are interested in potential avenues for improving these methods.

## Results

### Outline of the Challenge

Both rounds of the IronTract challenge followed the outline shown in Figure 1. *In vivo* tracer injections and *ex vivo* dMRI scanning were performed on two macaque brains. The dMRI data, acquired on a Cartesian grid, were resampled onto the two-shell of the HCP acquisition protocol^16^ (See *Methods for details*). We will refer to these datasets as diffusion spectrum imaging (*DSI*) and *HCP* respectively. The organizing team uploaded the data to the QMENTA platform (https://qmenta.com/irontract-challenge/) and the challenge teams could download them along with the tracer injection sites in the dMRI space. Each team analyzed the data with methods of their choice (*Methods, Analysis of dMRI data by challenge participants*). In round 1, this included image pre-processing, orientation reconstruction, tractography, and tractogram post-processing. In round 2, the pre- and post-processing steps were standardized across all teams. Each team produced tractograms with a range of thresholds and uploaded them to the QMENTA platform. A receiver operating characteristic (ROC) analysis was performed on the fly. The true positive rate (TPR) and false positive rate (FPR) were computed by voxel-wise comparison with the tracer. A partial area under the curve (AUC) was calculated, for which the maximum possible AUC score was 0.3 (*Methods, ROC analysis*). For the training case, participants were shown their score and were allowed to repeat data analysis and upload of results. Thus, participants tuned their analysis pipelines to maximize their score on the training case. Finally, they applied the optimized pipeline to the data from the validation case. The organizing team computed AUC scores on the validation case and used them for the final ranking of the challenge teams.

**Figure 1.**
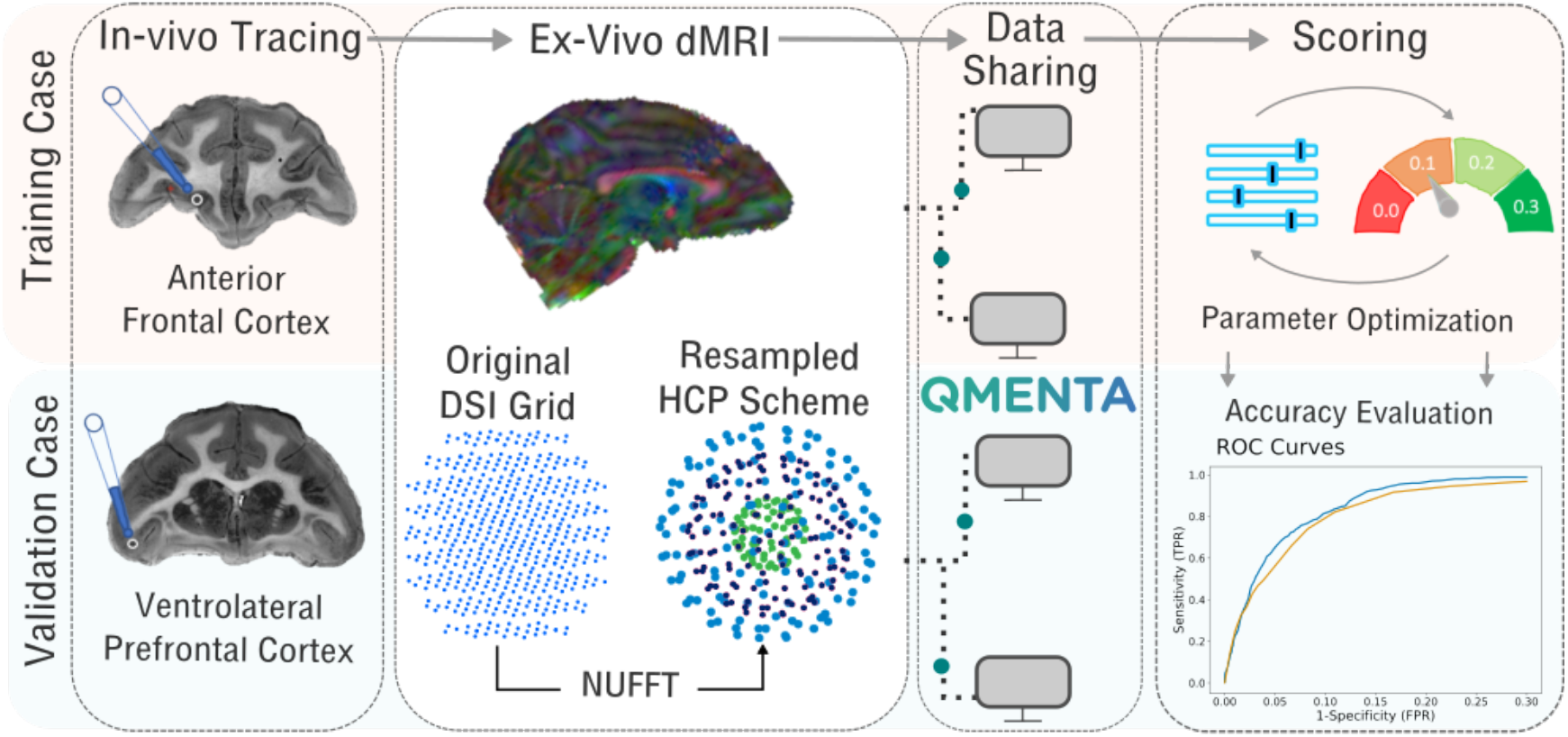
Overview of the IronTract Challenge. Data from two monkey brains, one with an injection in the anterior frontal cortex and one with an injection in the vlPFC, served as the training and validation case, respectively. *Ex vivo* dMRI data were acquired for both brains on a Cartesian grid (515 directions, bmax = 40,000 s/mm^2^) and resampled via NUFFT on the two shells of the HCP lifespan acquisition scheme, with b-values adjusted for *ex vivo* dMRI (93 directions with b=6000 s/mm^2^, 92 directions with b=12,000 s/mm^2^). Participants downloaded data and uploaded results on the QMENTA platform. For the training case, they received a score, allowing them to optimize their tractography pipeline. The optimized pipelines were then applied to the validation case for the final scores. In round 2, this procedure was repeated, but the pre- and post-processing (blue boxes) were standardized across teams.

### Round 1 results (variable pre- and post-processing)

Out of 30 registered teams, 12 completed the challenge (total submissions: 227; training: 186; validation: 38) and 16 final submissions were ranked. A detailed list is reported in Supplementary Table 1.

Overall, results from round 1 showed that, in both training and validation cases, no submission could achieve high TPR without also generating a large number of false positives (Figure 2A). Most submissions achieved TPRs higher than 0.8 only at FPRs higher than 0.2. Almost all submissions achieved higher accuracy in the training case (mean AUC=0.20) than in the validation case (mean AUC=0.16). Three teams only (Teams 1, 2, 6) obtained similar accuracy across datasets, with even higher accuracy for the validation case (Figure 2B). The AUC score of two of these three teams (Teams 1,2) was considerably higher (AUC > 0.23) than all other submissions (AUC ≤ 0.18) in the validation case. The overall highest score (AUC = 0.27) was obtained by Team 1, with a combination of the Robust and Unbiased Model-BAsed Spherical Deconvolution (Rumba-SD) method for orientation reconstruction^35^ and probabilistic tractography^36,37^ on the DSI data. Methods that used the DSI scheme achieved consistently high accuracy (Figure 2C, left), whereas methods that used the HCP scheme varied in their performance. However, the results suggest that, if analysis methods can be optimized carefully, the HCP acquisition may approach the accuracy of the much more demanding DSI acquisition. While most orientation reconstruction methods performed similarly in the training case, Rumba-SD^35^ outperformed the other submissions in the validation case (Figure 2C, center). Finally, probabilistic tractography approaches achieved overall higher accuracy scores (mean AUC = 0.20) than deterministic ones (mean AUC = 0.15), especially for the validation case (Figure 2C, right). (See Supplementary Figure 1 for performance by method.)

**Figure 2.**
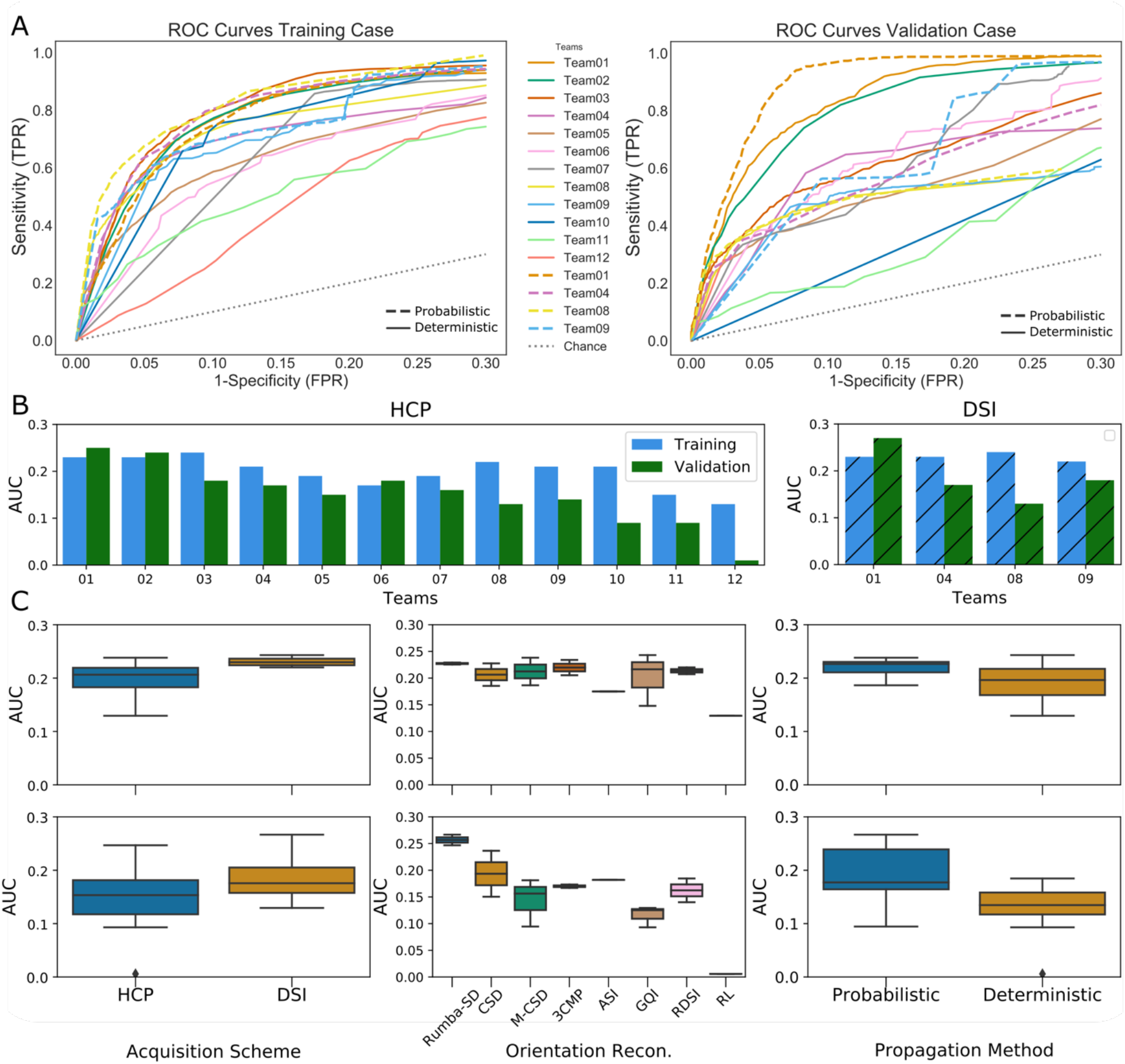
Round 1 results. **A)** ROC curves are shown for each submission. Results are shown for the training case (left) and validation case (right), and for the HCP (solid lines) and DSI (dashed line) acquisition schemes. **B)** Bar plots show the AUC score for each submission for the training case (blue) and validation (green) case, and for HCP and DSI sampling schemes. **C)** AUC scores are shown by acquisition scheme, orientation reconstruction method, and tractography propagation method for the training case (top) and the validation case (bottom). Rumba-SD = robust and unbiased model-based spherical deconvolution^35^; CSD= constrained spherical deconvolution^38^; M-CSD = multi-shell multi-tissue CSD^6,39^; 3Comp = three compartment model^40^; ASI = asymmetry spectrum imaging^41^; GQI = generalized Q-ball imaging^42^; RL= Richardson Lucy^43^, RDSI = radial diffusion spectrum imaging^44,45^.

### Sensitivity varies across white matter regions

We investigated how many of the white matter regions included in the tracer mask were correctly labeled by each Submission. To this end, every voxel included in the tracer mask in dMRI space was labeled by AY and CM. For the training case, voxels were assigned to one of 8 classes: anterior frontal white matter (AF); anterior limb of the internal capsule (ALIC); cingulum bundle (CB); corpus callosum (CC); external capsule (EC); medial prefrontal white matter (MPF); lateral prefrontal white matter (LPF); uncinate fasciculus (UF). For the validation case, voxels were assigned to one of 10 classes: ALIC; brainstem fibers (BS); commissural fibers (CF); CB; CC; EC; extreme capsule (EmC); LPF; thalamic fibers (ThF); UF. Figure 3 shows the TPR of each submission at the same specificity level (FPR=0.1) for each of these regions of interest (ROIs). Sensitivity was variable across regions, with similar patterns across submissions. In the training case, most teams labeled the EC, CC, and MPF correctly, but could reach the UF and CB only partially (Figure 3A).

**Figure 3.**
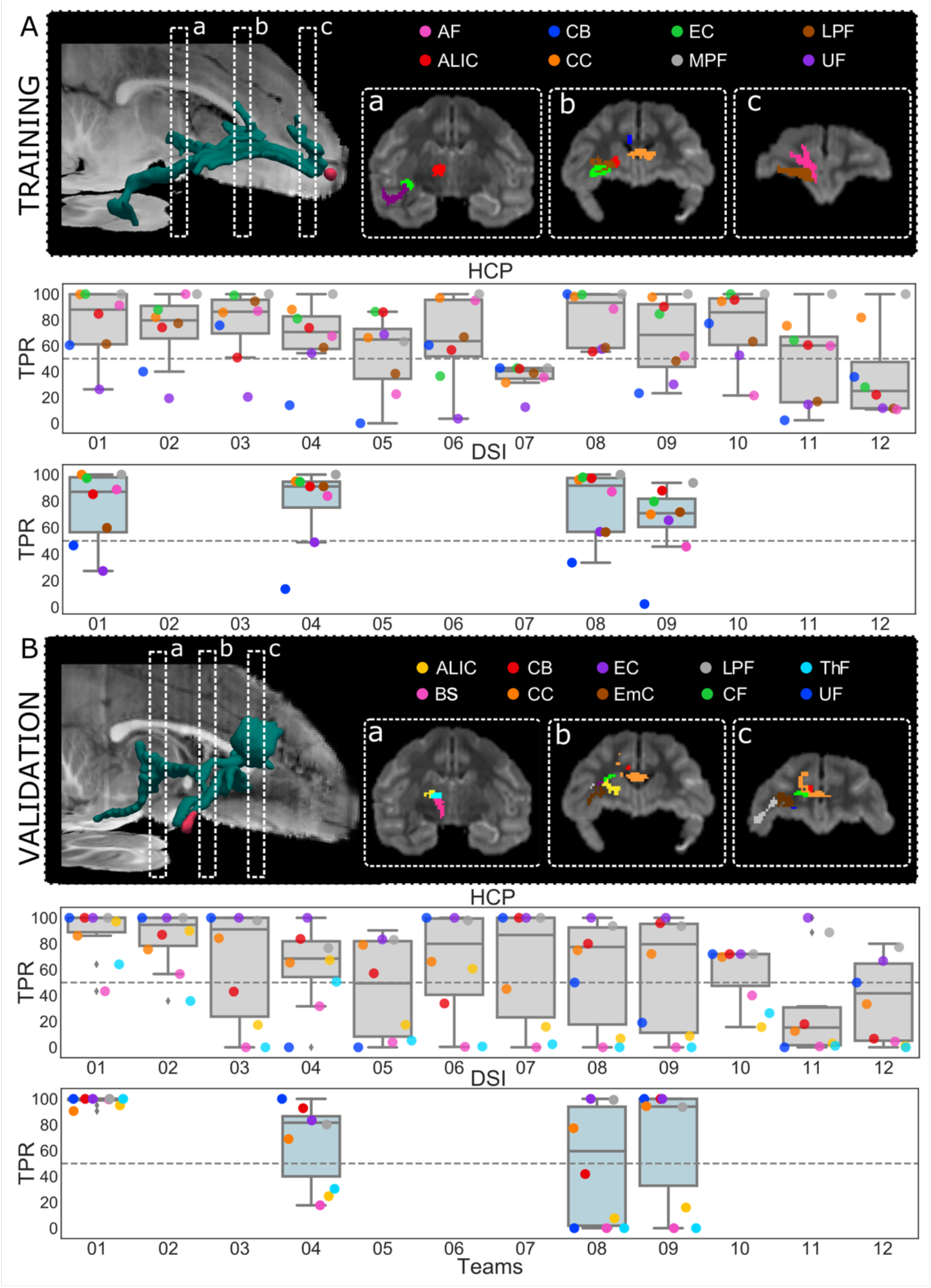
Performance by white-matter region. **A)** 3D rendering of the tracer mask for the training case, showing the location of the coronal slices that are displayed in boxes a, b, and c. The boxes show the main white-matter pathways present in the tracing. Boxplots overlaid with scatterplots show the TPR in each bundle for each submission, with the HCP scheme (top, light red) and the DSI scheme (bottom, light blue). **B)** The same results are presented for the validation case. All TPRs were evaluated at FPR=0.1. (AF = anterior frontal white matter; ALIC=anterior limb of the internal capsule; BS= brainstem fibers; CB = cingulum bundle; CC = corpus callosum; CF = commissural fibers; EC = external capsule; EmC = extreme capsule; LPF = lateral pre-frontal white matter; MPF = medial pre-frontal white matter; OF = orbitofrontal white matter; ThF = thalamic fibers; UF = uncinate fasciculus).

In the validation case, almost all methods could label the UF, EC, and LPF correctly but most of the submissions failed to reach regions located at a greater distance from the injection site, like the BF, ThF, and ALIC. In the training case several teams achieved similar performance as Team 1. In the validation case, however, where fine-tuning with respect to the ground truth was not possible, the performance of most teams deteriorated. The best result was achieved by the Rumba-SD model^35^ and probabilistic tractography^36,37^ on the DSI data (Team 1), which achieved a TPR higher than 0.9 for all the regions. There were clear differences in the bundles where errors occurred in the training *vs*. the validation case. This has to do with the fact that fibers starting from the two different injection sites enter these bundles from different angles (*See Localization of challenging areas)*.

### Round 2 results (standardized pre- and post-processing)

In round 2 the pre- and post-processing steps were standardized across teams. Participants downloaded pre-processed dMRI data from the QMENTA platform and were provided scripts to replicate the post-processing strategies that had been used by the two teams that had consistently good performance across both training and validation cases in round 1. These were: *(i) a Gaussian filtering strategy*, implemented by Team 1 in round 1; *(ii) an anatomical ROI* strategy, implemented by Team 2 in round 1 (see Methods for details).

Fourteen teams completed round 2 (259 total submission. Training: 105. Validation: 154). Of these, eleven also completed round 1, one completed round 1 but submitted results with a different pipeline in round 2, and two teams were new (Team 13 and Team 14). Some of the teams that had completed round 1 submitted results with new methods, in addition to regenerating results with the methods that they had used in round 1 but with the standardized pre- and post-processing. Fifty final submissions were ranked (Supplementary Table 2).

Results show that the performance of most returning teams improved when compared to round 1, as a result of applying the harmonized pre- and post-processing strategies. This improvement was greater for the validation case (2%-85%) than the training case (2%-30%) (Figure 4A). As a result, the difference in AUC score between the training and validation case decreased substantially in round 2 (Figure 4B). This led to many more teams achieving more similar performance between the training and validation case (Supplementary Figure 2). At the same FPR = 0.1, all submissions achieved higher TPR than in round 1 (Supplementary Figure 3).

**Figure 4.**
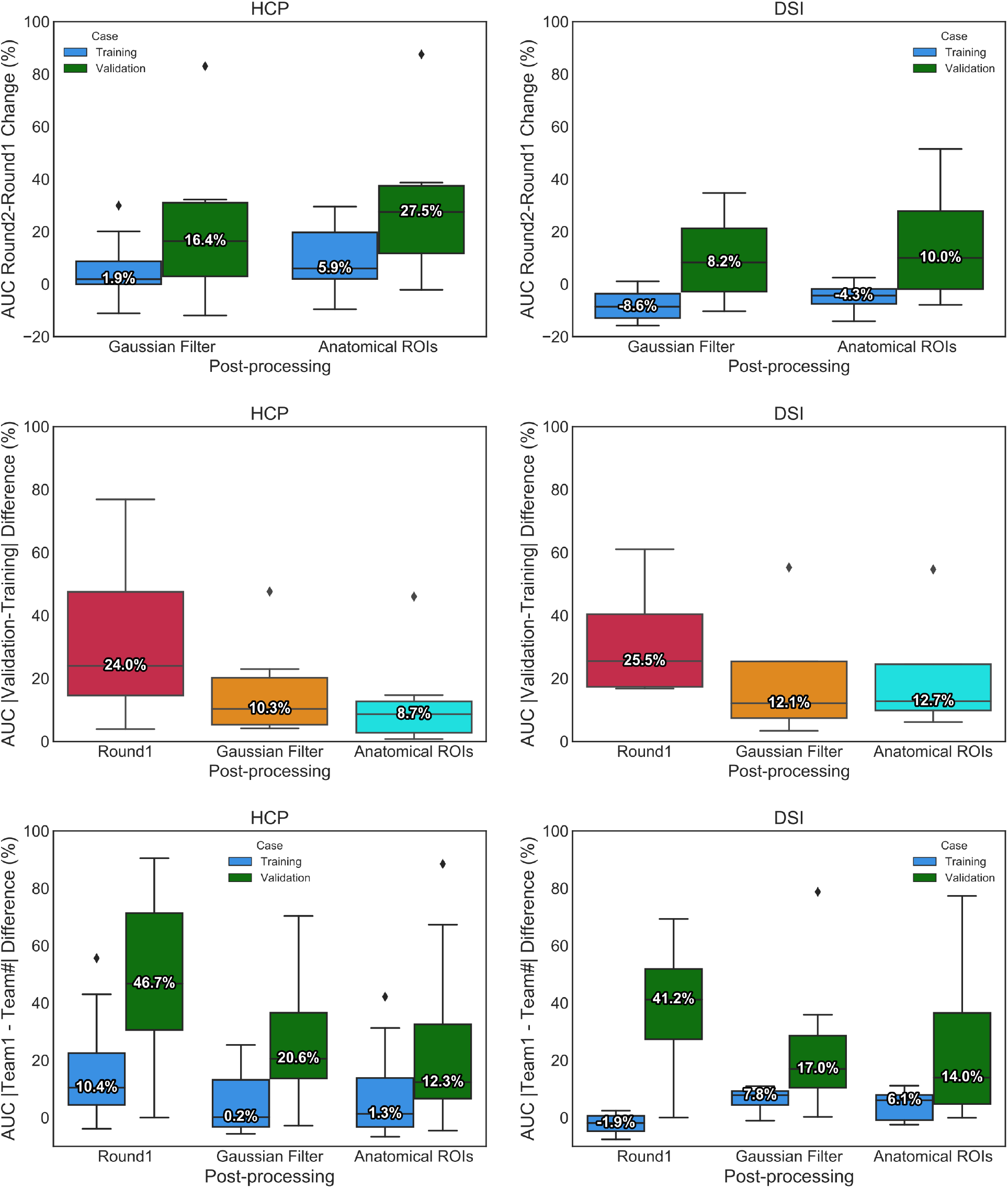
Effect of harmonized pre- and post-processing. **A)** Boxplots show the percent change in AUC scores between round 1 and round 2 for both post-processing strategies (Gaussian filter and anatomical ROIs). Results are shown for the training case (blue) and validation case (green), and for the HCP (left) and DSI (right) acquisition schemes. **B)** Difference in AUC scores between the training and validation cases, for round 1 and for each of the two post-processing strategies in round 2 *(Gaussian Filter* and *Anatomical ROIs*). **C)** Difference in AUC scores between each submission and the score achieved in round 1 by Team 1 for round 1 and the two post-processing strategies in round 2. Median percent change is indicated by a horizontal line in each plot.

Remarkably, post-processing by Gaussian filtering, which does not assume any prior anatomical knowledge, also improved results for most submissions (Figure 4B), leading to a training-validation percent difference only slightly higher than the one obtained when using the anatomical ROIs. Only two teams (Team 6 and Team 8) did not show improvement with Gaussian filtering and one of them (Team 8) did not show improvement with anatomical ROIs. These improvements allowed most teams to obtain higher scores, reducing the difference between their performance and that of Team 1, especially for the validation case (Figure 4C).

### Branching and turning fiber configurations are challenging for tractography

Figure 8 shows histograms of the number of teams that achieved a true positive (TP) (*i*.*e*., voxels included both in the tractogram and in the tracer mask) in each voxel of the tracer mask, at FPR = 0.1 (*Methods, Localization of challenging areas*).

These histograms are shown for round 1 and for each of the post-processing strategies adopted in round 2. The pre- and post-processing used in round 2 improved the overall coverage of the tracer masks by tractography. In the training case, the ALIC, CB, EC were labeled correctly by most teams (Figure 5, top, light blue arrows), while only few teams could label these regions in Round 1. The region where fibers turn sharply towards the temporal terminations of the UF remained challenging for all teams in both rounds (Figure 5, top, violet arrow). In the validation case, the biggest improvement was located where fibers coming from the ALIC branch into fibers entering the thalamus and fibers entering a narrow bundle of axons projecting down the brainstem. In round 2, more submissions labeled the thalamic fibers correctly and achieved improved coverage of the inferior brainstem fibers. Despite this improvement, this region continues to pose challenges for most teams (Figure 5, bottom, violet arrow). Like the UF region, this branch point is located further away from the injection/seed point than other regions in the tracing mask. Therefore, tractography needs to traverse other branching and turning points to get there and, as errors accumulate, the number of streamlines that reach these regions is small.

**Figure 5.**
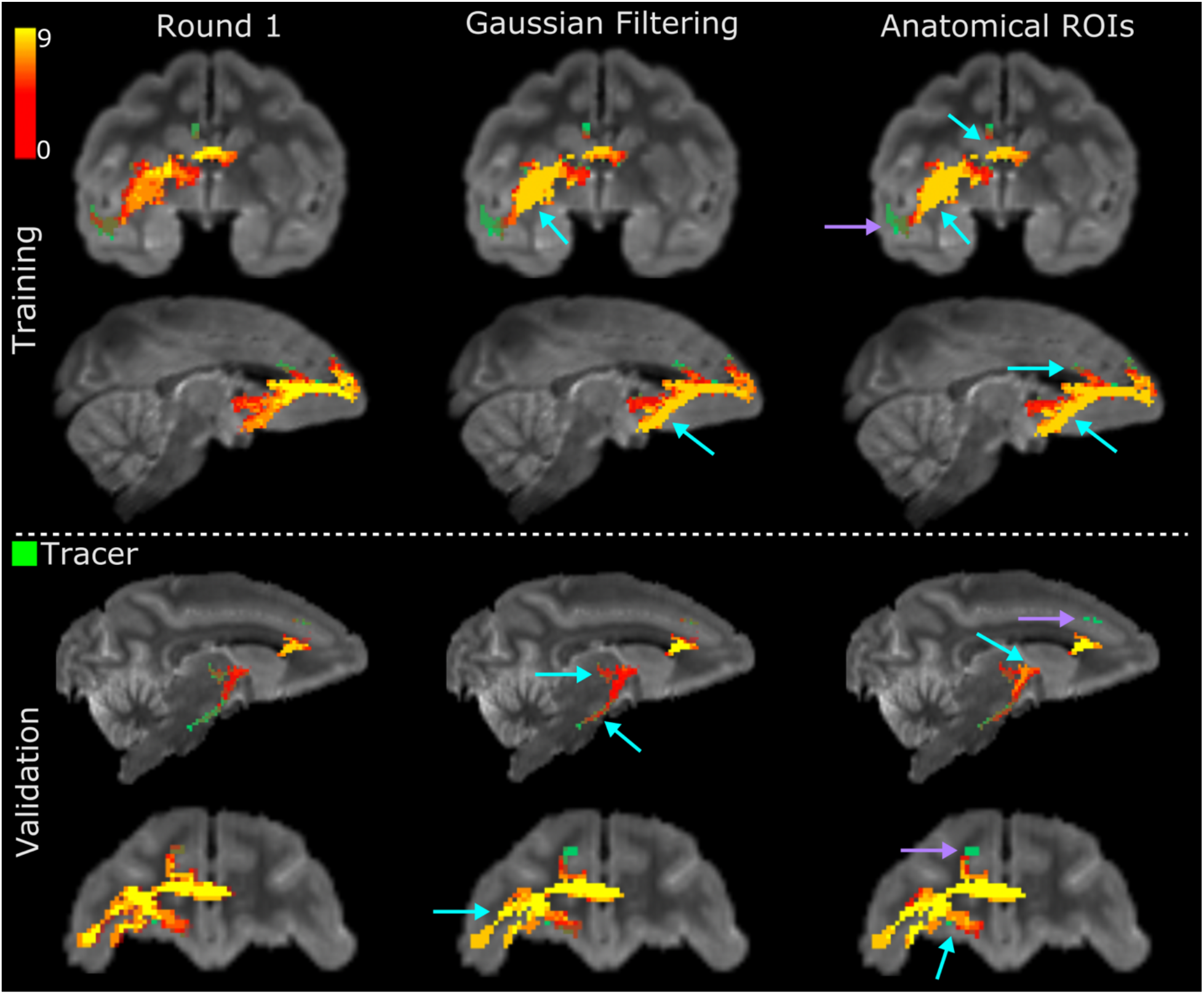
Number of teams reaching each voxel in the tracer mask. The heat maps are maximum intensity projections of the histograms of TPs across teams at FPR = 0.1, for the HCP acquisition scheme. The tracer mask is shown in green, under the heat maps. Results are shown for the training and validation case, and for round 1 and the two filtering strategies (Gaussian filtering, anatomical ROIs) in round 2. Only submissions that completed both rounds were included. Cyan arrows point to regions where the standardized pre- and post-processing round 2 led to improvement with respect to round 1. Violet arrows point to regions that remained challenging in both rounds.

We can better understand the nature of these errors by examining the false positives (FPs) that occur around these challenging areas, *i*.*e*., the paths that tractography chose to follow instead of the correct bundles. We identified two regions for the training case (UF and LPF) and two for the validation case (ALIC and EC) where the tracer and tractography trajectories consistently diverged in most submissions (Figure 6). We observed that in areas where fibers branch into two bundles, tractography tends to follow the least curved of the two and miss the other. Similarly, in areas where fibers take a sharp turn but, at the resolution of the dMRI data, overlap with a separate, less curved pathway, tractography follows the latter, instead of taking the turn. An example of such configuration is the area where the fibers coming from the EC turn towards the UF and the ILF (Figure 6B). Here tractography follows the ILF erroneously and fails to reach the UF terminations in the temporal lobe.

**Figure 6.**
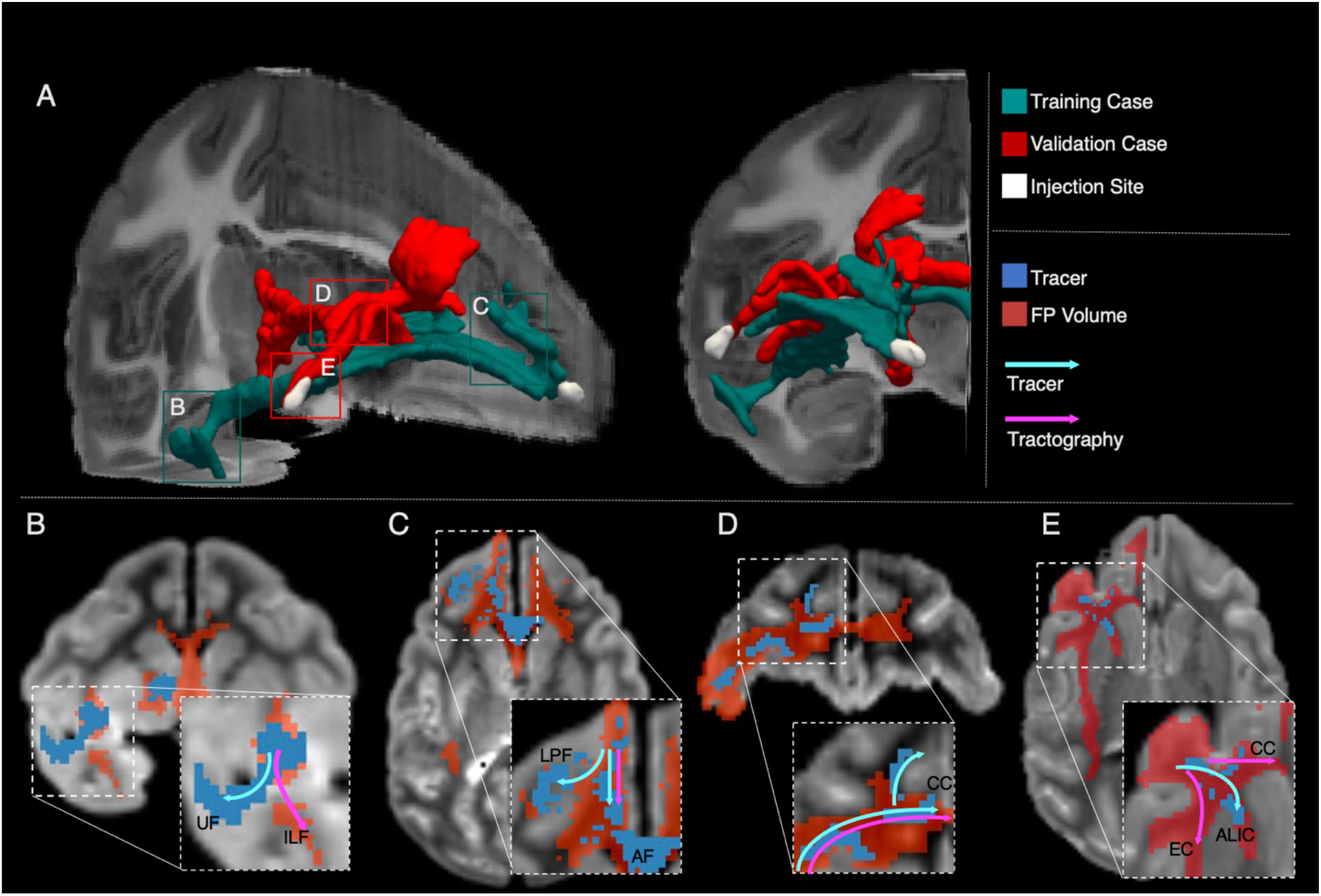
Challenging areas for tractography. **A)** 3D rendering of the tracer and injection site for the training (green) and validation (red) cases. Labeled boxes show the location of 2D views presented in B-E. **B**,**C)** A map of FPs is shown for one representative submission at FPR = 0.1 (red), overlaid by the tracer mask (blue) for the training case. Streamlines follow the ILF, instead of turning into the UF (B). Streamlines continue into the AF instead of fanning into the LPF (C). **D**,**E)** A map of FPs is shown for one representative submission at FPR = 0.1 (red), overlaid by the tracer mask (blue) for the validation case. Streamlines continue in the body of the CC to project contralaterally and miss the turn into the superior frontal gyrus (D). Tractography follows paths of lower curvature in the body of the CC and in the EC, instead of projecting into the ALIC (E). AF: antero-frontal white matter; ALIC: anterior limb of the internal capsule; CC: corpus callosum; EC: external capsule; ILF: inferior longitudinal fasciculus; LPF: lateral pre-frontal white matter; UF: uncinate fasciculus.

Fanning regions also lead to errors in tractography. In the training case, fibers exiting the injection site branch from the main bundle, which is sometimes referred to as the “stalk”, and fan out towards the dorsolateral prefrontal cortex. Here tractography follows the main stalk, continuing in the frontal white matter and does not turn supero-lateral to then fan into the LPF (Figure 6C). In the validation case, most teams showed false negatives (FNs), *i*.*e*., voxels included in the tracer mask but not in the tractogram, in the supero-frontal projections of the CC (Figure 5). Figure 6D shows that here tractography continues into the body of the CC to project to contralateral areas, missing the sharp turn of CC projections to the superior frontal gyrus. Another region of the validation case that showed significant FNs across submissions was the region where fibers enter the ALIC. Here tractography prefers following the direction of least curvature in the CC body and into the big bundle of anterior-posterior fibers stemming from the EC, rather than turning into the smaller ALIC (Figure 6E).

### Sharper diffusion profiles do not always lead to more accurate tractography

We compared the orientation distribution functions (ODFs) from different submissions in an area that was consistently challenging across methods. This was where fibers branched into thalamic and brainstem fibers (Figure 7) (*Methods, Comparison of orientation distribution functions*). All submissions identified two fiber populations correctly in the superior part of this region, where fibers branched, and one main fiber population in the inferior part, where fibers projected caudally to the brainstem. However, there were differences in the sharpness of the ODFs. Interestingly, the submissions that achieved the highest accuracy had somewhat less sharp diffusion profiles in the superior part of the ROI, where the two sets of fibers diverge (Figure 7B-D). However, one of the submissions achieving lower accuracy also shows somewhat less sharp ODFs (Figure 7O).

**Figure 7.**
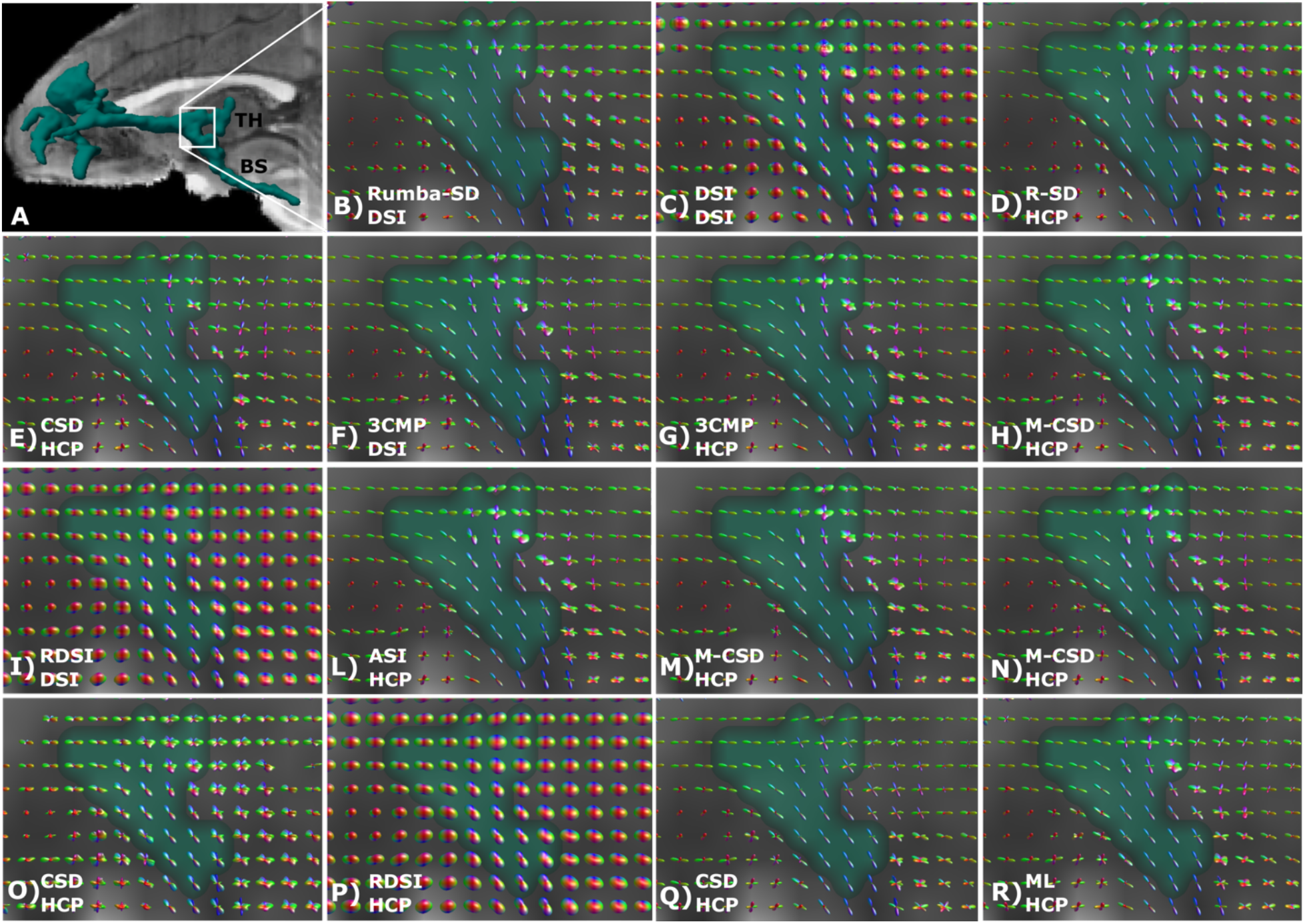
Comparison of ODFs across submissions. **A)** 3D rendering of the tracer mask from the validation case, showing the location of the magnification region where thalamic (TH) and brainstem (BS) fibers branch. **B-R)** ODFs for each submission are visualized for the region shown in A. Submissions are ordered based on the AUC score obtained for the validation case in round 2. ASI: asymmetry spectrum imaging^41^; 3CMP: three compartment model^40^; CSD: constrained spherical deconvolution^38^; DSI: Diffusion spectrum imaging^46^; M-CSD: multi-shell multi-tissue CSD^6,39^; ML: machine learning-based reconstruction^47^; RDSI: radial diffusion spectrum imaging^44,45^; Rumba-SD: robust and unbiased model-based spherical deconvolution^35^.

We quantified the sharpness of the ODFs by computing the dispersion of each peak in each voxel. Figure 8 shows plots of the average dispersion in seven ROIs from the training and validation case. We selected both regions with complex fiber configurations (UF, CB, CC, EC-IC, TH-BS) and regions that should mainly contain single fiber orientations, like the body of the CC (CCb) and BS. Figure 8 shows that, although ODF dispersion was not the only factor that determined accuracy, submissions that achieved higher AUC scores had less sharp ODFs, especially in regions with turning, fanning, and branching fiber configurations (TH-BS, CC, UF).

**Figure 8.**
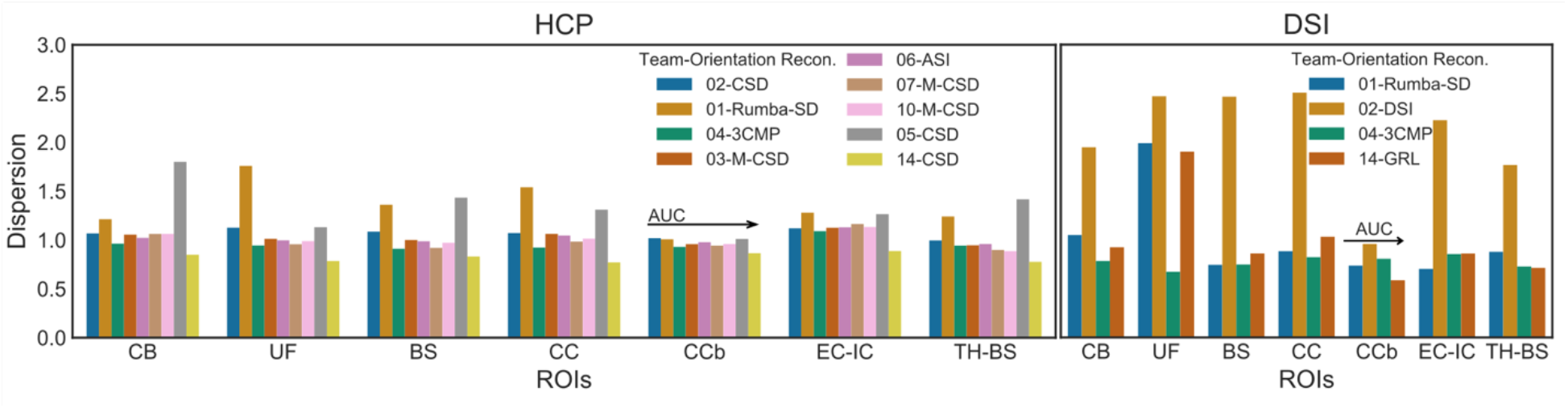
Effect of ODF dispersion and peak orientation on the accuracy of tractography. Bar plots of mean dispersion for each submission and each sampling scheme across different ROIs from the training and validation cases. For each ROI, teams are ordered along the x-axis based on AUC score for the validation case in round 2. Note that, as dispersion affects only methods that sample orientations from ODF, we excluded methods that follow the peak orientation exclusively. Training case: CB = cingulum bundle, UF = uncinate fasciculus. Validation case: BS = brainstem, CC = corpus callosum, CCb = body of the corpus callosum, EC-IC = external capsule – internal capsule, TH-BS = thalamus – brainstem. ASI = asymmetry spectrum imaging^41^; 3CMP = three compartment model^40^; CSD = constrained spherical deconvolution^38^; DSI = Diffusion spectrum imaging^46^;; M-CSD = multi-shell multi-tissue CSD^39^; GRL= generalized Richardson-Lucy^48^; ML = machine learning-based reconstruction^47^; RDSI = radial diffusion spectrum imaging^44,45^; Rumba-SD = robust and unbiased model-based spherical deconvolution^35^.

## Discussion

The IronTract Challenge evaluated a variety of state-of-the-art tractography methods on high-angular and spatial resolution dMRI data by quantitative voxel-wise comparison to anatomic tracing data in the same NHP brains. This effort differed from previous tractography challenges in several ways. First, the dMRI acquisition protocol allowed us to evaluate HCP-style and DSI acquisition schemes in real brain data. Second, the availability of both dMRI and tracer data in the same brains allowed the precise localization of tractography errors and challenging fiber configurations. Third, a training and validation case with different injection sites allowed us to evaluate the robustness of submissions across seed areas. Fourth, a full ROC analysis allowed us to compare the sensitivity of different methods at the same level of specificity. Fifth, by iterating over the results in a second round, where all teams used the same pre- and post-processing steps, we disentangled the contribution of these steps from that of the orientation reconstruction and tractography steps. Our results provide insights into the optimal processing strategies for widely available, HCP-style data. They also reveal why errors occur even with these state-of-the-art acquisition and analysis techniques, thus pointing to possible areas of improvement for future methodological development.

### The effect of acquisition scheme and propagation method

We compared an HCP-style, two-shell acquisition scheme with a much more densely sampled DSI scheme. Overall, higher accuracy was achieved by methods that used the full DSI data (515 diffusion volumes) (Figure 2). However, a few of the methods that used the HCP data approached the accuracy of the DSI methods. For methods that could be applied to both schemes, the loss in accuracy when using HCP versus DSI data was lower than 10% (Supplementary Table 1, Figure 2). This illustrates that when analysis methods are carefully optimized, the two-shell HCP scheme represents an advantageous trade-off between accuracy and acquisition time, given that DSI acquisition involves 2.8 times more directions and 3.3 times higher maximum b-value. Previous validation studies showed that DSI produces more accurate fiber orientation estimates both in simulations^49^ and in comparison to optical imaging measurements^50^. In this study, the most accurate submission was obtained using DSI data. While a full DSI acquisition is time-consuming, compressed sensing (CS) allows DSI data to be reconstructed from undersampled q-space^4,51,52^. A recent post mortem validation study showed that a CS-DSI protocol with 171 directions (similar to the number of directions in the two-shell HCP protocol), preserves the high angular accuracy of fully sampled DSI^53^. Thus, it is a viable alternative that combines the benefits of shell and grid acquisitions.

In regard to the propagation method, we found that probabilistic tractography led to overall higher AUC (mean AUC: 0.22) than deterministic tractography (mean AUC = 0.17). This was particularly true for the validation case, where pipelines were not optimized with respect to the ground truth (Figure 2A, 2C). This confirms the overall lower sensitivity of deterministic approaches at the same level of specificity^28,29^. Probabilistic tractography led to better bundle coverage (Supplementary Figure 1,4). Three deterministic submissions could not reach all the bundles labeled in the validation case, and the other ones did so at a much higher FPR than the probabilistic methods (Supplementary Figure 4). This was especially true for white matter regions located further away from the injection site/seed (Figure 3B).

### The effect of orientation reconstruction method

Differences between the ODFs from the various submissions were mostly subtle. Our results suggest that there is no simple, one-to-one mapping between ODF characteristics and the accuracy of tractography (Figure 8, Supplementary Figure 7). This result is in line with a recent study that found that there is no single optimal method for all different fiber configurations^54^.

However, the dispersion of the ODFs does seem to play a role. The conventional wisdom is that sharper ODFs are better because they help resolve crossing fibers with small inter-fiber angles^54^. However, the ODFs from the winning method (Rumba-SD) showed higher dispersion than ODFs from most of the other submissions. This was the case in almost all selected ROIs and especially in those that included branching, fanning, or turning fibers (Figure 8). Less sharp ODFs, when combined with probabilistic tractography, allow a broader range of orientations to be sampled from the same ODF peak. This can be beneficial in areas of branching or fanning. Areas where fibers take sharp turns remain a challenge for all methods. They can only be resolved by relaxing bending angle thresholds to a degree where the FPR becomes prohibitively high.

In a previous study, we evaluated a different set of tractography methods on the dataset that we refer to as the training set here^29^. We observed the highest accuracy from the combination of probabilistic tractography with GQI, a reconstruction method that does not produce particularly sharp ODFs. The performance of probabilistic GQI in that study (TPR < 0.7 at FPR = 0.1) was lower than the performance of probabilistic Rumba-SD in the present study (TPR = 0.74 at FPR = 0.1). However, it may be worth revisiting the probabilistic GQI approach with the optimized pre- and post-processing methods of the IronTract Challenge.

### The effect of pre- and post-processing

In the second round of the challenge, we investigated the extent to which the pre- and post-processing strategies had contributed to the higher robustness achieved by teams 1 and 2 in the first round. When the remaining teams used the same strategies, accuracy improved for almost all submissions (Figure 4). This improvement was higher for the validation than the training case, *i*.*e*., the accuracy of tractography became more robust to the location of the seed region. More specifically, accuracy improved in some regions that proved challenging in round 1 (Figure 5).

While we did not study the effects of the pre- and post-processing separately, prior work studied the effects of some pre-processing steps on the accuracy of diffusion orientation estimates^49^. They found that denoising improved orientation accuracy up to 30%–40%. Approximately half of the teams had applied denoising in round 1 and only four teams had performed eddy-current correction. These steps were included in the standardized pre-processing of round 2.

The improved accuracy obtained with the use of *a priori* anatomical ROIs was expected. The more surprising result was that post-processing with a simple Gaussian filter, which requires no prior anatomical information, increased the AUC by up to 80%, a benefit similar to the use of anatomical ROIs (Supplementary Figure 2). While harmonizing pre- and post-processing in round 2 decreased the difference in AUC score between all the submissions and Team 1, the latter continued to achieve the highest accuracy. When using DSI data, Team 1 could reach a much higher TPR than all other submissions (TPR = 0.96 at FPR=0.1), suggesting that its pre- and post-processing strategies were not the only factors contributing to its high performance.

### Localization of challenging areas

Having data from anatomic tracing and dMRI in the same monkey brain allowed us to identify the regions where tractography errors occurred consistently across submissions. These included regions where fibers branched into smaller bundles, or where they took a sharp turn to enter a bundle (Figures 5, 6). These results agree with previous validation studies^21,29^ and illustrate the importance of anatomic tracing for identifying realistic failure modes of tractography that go beyond the simple crossing fiber configurations used in digital or physical phantoms. Almost all submissions were successful in identifying projections that ran through major crossing regions (Figures 3, 5). However, many methods had trouble following fibers that branched into smaller bundles or fanned off the main bundle (Figure 5, 6). These results highlight the need for further validation and development of tractography methods that go beyond the crossing-fiber paradigm.

### Robustness across seed areas

Our training and validation cases allowed us to evaluate the robustness of tractography methods across different seed areas. The two injection sites, while projecting through similar white-matter pathways (Figure 3), follow very different routes to reach these pathways and pose different challenges to tractography. In the training case, the injection site is in the frontal pole. From here, most fibers travel straight posteriorly to enter the internal and external capsule. The most challenging areas are where fibers fan out into the LPF or turn into the UF and CB (Figure 5,6). In the validation case, the injection site is in the vlPFC. From here, fibers need to first course medially and take a more complicated and curved trajectory before entering the capsules. The ALIC shows lower TPs in the validation case than in the training case (Figure 3), and the most challenging area is located posterior to the ALIC where fibers branch into thalamic and brainstem fibers (Figure 5).

For most of the submissions, optimizing the methods with respect to accuracy for one seed/injection region did not guarantee optimal performance for another region, with a 25% average decrease in AUC score between the training and the validation case (Figure 2). Only two teams could achieve high accuracy for both injection sites. One of these two teams used anatomical ROIs, based on general knowledge on the connections of the prefrontal cortex from previous tracer experiments^55^, illustrating the importance of such experiments for mapping the organizational rules of white matter projections. In future studies, we intend to investigate a wider variety of injection sites and evaluate whether these conclusions generalize to different brain areas.

### Optimal data processing for the HCP protocol

One of the main goals of the IronTract challenge was to identify optimal processing strategies for the widely used, two-shell HCP acquisition scheme. Our results can inform various methodological choices that have to be made when analyzing such data, including pre-processing, orientation reconstruction, tractography, post-processing, and thresholding. When these choices were made as summarized below, tractography reconstructed 8 out of the 10 bundles present in the tracer mask with FPR = 0.05, and it reconstructed all 10 with FPR = 0.1 (Supplementary Figure 4).

#### Pre-processing

The winning pipeline included denoising^56^, corrections for Gibbs ringing^57^, and motion/eddy-current distortions^58,59^, all sensible and widely used procedures.

#### Orientation reconstruction

The method that achieved the highest performance was Rumba-SD (Figures 3, Supplementary Figures 4,6,8). Its estimation framework relies on Rician and noncentral Chi likelihood models, which accommodate realistic MRI noise, and a 3D total-variation spatial regularization term, which promotes continuity and smoothness along individual tracts by taking into account the spatial correlation among adjacent voxels^35^. While this is a relatively newer method, we note that high accuracy and robustness were also achieved by classical reconstruction methods like CSD^38^ (applied on the high-b shell only) and DSI^46^. However, these results were specific to Team 2, who supplemented these methods with anatomical ROIs. The 3CMP^40^ and M-CSD^6,39^ also achieved relatively higher accuracy and lower reconstruction error than other methods (Figures 5, 6).

#### Tractography

Our results concur with previous studies that showed the higher sensitivity of probabilistic methods, when compared to their deterministic counterparts at the same specificity^28,29,60^.

#### Post-processing

Simple Gaussian post-filtering improved the accuracy of most tractography methods used in this challenge, as well as their robustness to the location of the seed region. The use of inclusion ROIs based on prior anatomical knowledge led to small additional gains in performance.

#### Thresholding

Most methods required a rather low threshold (< 2% of the maximum value of the tractogram) to reach all the main bundles present in the tracer (Supplementary Figure 5). This is in agreement with a prior finding that the biggest changes in tractograms occur between thresholds of approximately 2–3%, above which the sensitivity of tractography decreases dramatically^21^. We note that we focused on optimal thresholds for reconstructing all the bundles that the injection site projects to, which is a task that requires high sensitivity. In other tasks, such as constructing whole-brain connectivity matrices, high specificity may be more important. In that case, one may want to use more stringent thresholds and accept that only a subset of the true connections will be included.

### Limitations

The main limitation of using tracer injections to validate dMRI tractography is that such studies cannot be performed in the human brain. Human and NHP brains differ in terms of both absolute and relative sizes of different gray and white-matter structures. However, similarities in position, cytoarchitectonics, connections, and behavior indicate that the overall organization of brain circuitry is relatively comparable^61,62^. In particular, the relative positions of different brain regions, as well as the obstacles the fibers encounter on their way from one area to another, are comparable. As a result, similar fiber geometries (crossing, branching, turning, fanning) exist in similar locations of the NHP and human brain. Thus, important insights can be gained from the performance of tractography methods in NHP brains.

The present study was limited to two injection/seed areas. Furthermore, we used binary tracer and tractography maps, *i*.*e*., we only compared the presence or absence of labeled axons and tractography streamlines at each voxel, rather than their density. Automated methods for segmenting and quantifying the tracer maps will be critical for extending these analyses in the future.

Other limitations of tracer validation studies include imperfect tracer uptake or imperfect alignment of histology and dMRI data. The injections used in this study passed rigorous quality assurance checks at Dr. Haber’s laboratory and had high-quality transport. The manual annotation of the axon bundles and their alignment to the dMRI volumes were also checked by Dr. Haber and refined at multiple stages.

## Conclusion

As part of the IronTract challenge we undertook a comprehensive, quantitative, voxel-wise assessment of tractography accuracy across different tractography pipelines, acquisition schemes, and seed areas. This allowed us to identify common failure modes of tractography for both established and new tractography algorithms and to propose optimized strategies for analyzing dMRI data that have been acquired with state-of-the-art, high angular resolution techniques, including the popular two-shell acquisition scheme employed by the lifespan and disease HCP. The IronTract Challenge remains open (https://qmenta.com/irontract-challenge/) and we plan to expand its scope in future iterations. We hope that it can serve as a valuable validation tool for both users and developers of dMRI analysis methods.

## Methods

### Data description

The training and validation cases used in this challenge are part of a previously described dataset that consists of *in vivo* tracing and high-resolution *ex vivo* dMRI acquired in the same macaque brains^29–31^.

### Tracer injections

The training and validation datasets came from two different male rhesus macaques. The former received an injection of the anterograde/bidirectional tracer Lucifer Yellow in the anterior frontal cortex (frontal pole). The latter received an injection of the anterograde/bidirectional tracer Fluorescein in the ventrolateral prefrontal cortex (vlPFC). Surgery and tissue preparation were performed at the University of Rochester Medical Center. Details of these procedures were described previously^30,55,63^. Briefly, each monkey received an injection of a bidirectional tracer conjugated with dextran amine (40–50 nl, 10% in 0.1 M phosphate buffer, pH 7.4; Invitrogen). Twelve days after the injection, animals were perfused and their brains were postfixed overnight and cryoprotected in increasing gradients of sucrose (10, 20, and 30%). All experiments were performed in accordance with the Institute of Laboratory Animal Resources Guide for the Care and Use of Laboratory Animals and approved by the University of Rochester Committee on Animal Resources.

### dMRI data acquisition

After fixation, the brains were scanned in a small-bore 4.7T Bruker BioSpin scanner (maximum gradient strength 480 mT/m) using a 3D EPI sequence with the following parameters: TR = 750 ms, TE = 43 ms, δ = 15 ms, Δ = 19 ms, maximum b = 40,000 s/mm^2^, matrix size 96 × 96 × 112, 0.7 mm isotropic resolution. Brains were submerged in liquid Fomblin to eliminate susceptibility artifacts. We acquired 1 non-diffusion weighted (b = 0 s/mm^2^) volume and 514 diffusion-weighted volumes corresponding to a Cartesian lattice in q-space. The total acquisition time was 48 hours. We refer to this q-space sampling scheme data as diffusion spectrum imaging (DSI).

We resampled the data onto q-shells, following a methodology that was previously described and validated^50,53^. It involves approximating data points distributed on spheres in q-space from data points distributed on a Cartesian grid, using a fast implementation of the non-uniform fast Fourier transform (NUFFT)^64^. We followed this procedure to generate data on the two q-shells of the lifespan and disease HCP acquisition protocol^16^. This *in vivo* protocol includes 93 directions with b=1500 and 92 directions with 3000 s/mm^2^. We multiplied these b-values by the 4x factor required to achieve comparable diffusion contrast *ex vivo* as *in vivo*^65^, *i*.*e*., we used b=6000 and 12000 s/mm^2^. We refer to this q-space sampling scheme as HCP.

### Histological processing

Following whole-brain *ex vivo* dMRI, the brains were returned to the University of Rochester for histological processing. They were sectioned in 50 μm thick coronal slices on a freezing microtome into 0.1 m phosphate buffer or cryoprotectant solution as previously described^66^. An undistorted photo of the blockface was taken before cutting for use in image registration (See *Registration of tracer and dMRI data*). Immunocytochemistry was then performed on every 8th slice to visualize the transported tracer, resulting in an inter-slice resolution of 400 µm. Additional details on the histological procedures can be found elsewhere^55,67,68^. Labeled fiber bundles were outlined under dark-field illumination with a 4.0 or 6.4x objective, using Neurolucida software (MBF Bioscience). Fibers traveling together were outlined as a group or bundle. Axons were charted as they left the tracer injection site and followed through the right hemisphere, until the anterior commissure. The 2D outlines were combined across slices using IMOD software (Boulder Laboratory^69^) to create 3D renderings of the structures and pathways as they traveled through them. These were used to further refine bundle contours and ensure spatial consistency across sections.

### Registration of tracer and dMRI data

Each histology slice was registered to its corresponding blockface using a 2D robust affine registration^70^, followed by a 2D symmetric diffeomorphic registration^71^. Blockface images were then stacked to create a 3D volume and registered to the b=0 dMRI volume using a 3D affine registration followed by a 3D diffeomorphic registration, with the same methods as above. The computed transformations were then applied to the tracer mask and the injection site mask, to map them into dMRI space. The transformed injection site mask was shared with challenge participants, to be used as the seed region for tractography.

### Analysis of dMRI data by challenge participants in Round 1

In the first round, teams were provided raw dMRI data. They were allowed to use the q-space sampling scheme and analysis methods of their choice. A detailed description of the methods that each team used in this round, including pre-processing, orientation reconstruction method, tractography, and post-processing, are provided in the Supplementary note 1. Both probabilistic and deterministic tractography approaches were deployed, with a variety of orientation reconstruction methods. Participants were asked to generate tractograms at multiple thresholds by varying one or more parameters of their choice. The most common choices were lower thresholds on probability, for submissions that used a probabilistic tractography algorithm; and upper thresholds on the bending angle, sometimes combined with lower thresholds on fractional anisotropy or other microstructural parameters, for submissions that used a deterministic tractography algorithm.

For each submission, participants uploaded a series of tractograms, obtained with different thresholds, to the QMENTA platform. A score was computed on the fly by comparing the tractograms to the tracer data (see *ROC analysis*). For the training case, the platform generated a performance report, including the AUC score, and made it available to the participant. Participants could repeat their analysis, upload, and score any number of times, allowing them to fine-tune the free parameters of their methods and optimize their score. They then applied their optimized analysis pipeline to the dMRI data from the validation case and uploaded the resulting tractograms to the QMENTA platform. The final AUC scores were computed from the validation case and used to rank the teams.

### Analysis of dMRI data by challenge participants in Round 2

In the second round, analysis and scoring of the training and validation cases were performed as described above. The difference was that the pre- and post-processing steps were standardized across teams. Participants downloaded pre-processed dMRI data from the QMENTA platform and were provided scripts for the post-processing steps.

#### Pre-processing

This followed the dMRI pre-processing procedures that had been used in round 1 by Team 1, the team that achieved the best performance (*Results, Round 1 Results*). They included denoising^56^ and correction for Gibbs ringing^57^ in MRtrix3^72^, and correction for motion and eddy-current distortions in FSL^58,59^. A binary dilation was applied to the tracer injection seed/point.

#### Orientation reconstruction and tractography

Teams were asked to apply the same orientation reconstruction and tractography methods as in round 1, if they had participated in round 1, or any methods of their choice otherwise.

#### Post-processing

This replicated the post-processing strategies that had been used by the two teams that had consistently good performance across both training and validation cases in round 1 *(Results, Round 1 Results*). *(i) Gaussian filtering*. This strategy had been implemented by Team 1 in round 1. It included the application of a Gaussian filter with sigma = 0.5 to increase coverage, followed by an iterative thresholding of 200 steps on the log of the streamline count, for a total of 200 output tractogram volumes. *(ii) Anatomical ROIs*. This strategy had been implemented by Team 2 in round 1. ROIs from the PennCHOP macaque atlas^73^ were transformed to the space of each dMRI dataset. Only streamlines intersecting at least one of these ROIs were retained. The ROIs were selected on the base of general knowledge of projections of the prefrontal cortex^55^ and were located in: the cingulum bundle, the genu of the corpus callosum, the external capsule, the anterior limb of the internal capsule, and the uncinate fasciculus. For round 2, after applying the anatomical ROIs, the same smoothing (sigma = 0.5) and iterative thresholding (200 steps on the log of the streamline count) as in the Gaussian filtering strategy were performed.

### ROC analysis

We adopted the area under the ROC curve (AUC) as our main performance score (See *Supplementary Note 3* for additional metrics). The ROC analysis was performed as follows. For each of the submitted tractograms, we obtained the numbers of voxels that were true positive (TP; voxels included both in the tractogram and in the tracer mask), true negative (TN; voxels included neither in the tractogram nor in the tracer mask), false positive (FP; voxels included in the tractogram but not in the tracer mask), and false negative (FN; voxels included in the tracer mask but not in the tractogram). The true-positive rate (TPR) and false-positive rate (FPR) were then calculated as follows:

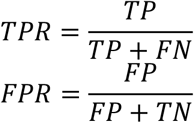

This was repeated for all tractograms in a submission, which had been thresholded at different levels (either with the thresholding method chosen by each team in round 1, or with the standardized thresholding method in round 2). We obtained the ROC curve of each submission by plotting the TPR as a function of FPR. We computed a partial AUC score, *i*.*e*., the area under the ROC curve for FPR in the [0,0.3] range. Thus, the maximum possible AUC score was 0.3. The choice of this range was based on prior results showing that deterministic tractography methods cannot always achieve FPRs outside this range^29^.

### Localization of challenging areas

Having tracer and dMRI data from the same brain allows us to identify the exact locations where tractography goes wrong, and thus the fiber geometries that are consistently challenging across tractography methods. To this end, we extracted a map of TP voxels at FPR = 0.1, for each of the submissions that participated in both rounds of the challenge. We binarized these maps and summed them across all submissions. This yielded a histogram that showed the number of teams that achieved a TP in each voxel of the tracer mask. This allowed us to identify the locations where errors occurred consistently across tractography methods in round 1, and to examine whether the pre- and post-processing steps that were applied in round 2 mitigated these common errors.

### Comparison of orientation distribution functions

After the end of the challenge, we asked participants to share the ODFs from their final submissions, to examine if the ODFs played a role in the performance differences between teams. All ODFs were projected onto a common set of 362 directions that were distributed uniformly on the half sphere. This direction set was generated by the electrostatic repulsion model^74^, as implemented in DIPY^36^. We then normalized the ODFs by the maximum ODF value and converted their amplitudes to their spherical harmonic representation in MRtrix3 (*lmax*=12)^72^. For each submission, we extracted a voxel-wise map of orientation dispersion by computing the mean dispersion of the ODF lobes inside the voxel^75^. We included only ODF lobes with peak amplitudes larger than 0.2 times the maximum ODF amplitude.

## Data Availability

The authors declare that the data supporting the findings of this study are available on the QMENTA platform (https://qmenta.com/irontract-challenge/). Detailed information on how to reproduce the tractograms generated by the Challenge Teams, and links to code repositories are provided in the Supplementary information.

## Acknowledgements

Data acquisition was supported by the National Institute of Mental Health (R01-MH045573, P50-MH106435). Additional research support was provided by the National Institute of Biomedical Imaging and Bioengineering (R01-EB021265) and the National Institute of Neurological Disorders and Stroke (R01-NS119911). Imaging was carried out at the Athinoula A. Martinos Center for Biomedical Imaging at the Massachusetts General Hospital, using resources provided by the Center for Functional Neuroimaging Technologies, P41-EB015896, a P41 Biotechnology Resource Grant, and instrumentation supported by the NIH Shared Instrumentation Grant Program (S10RR016811, S10RR023401, S10RR019307, and S10RR023043). Andrey Zhylka is supported by the European Union’s Horizon 2020 research and innovation program under the Marie Sklodowska-Curie grant (765148). Ye Wu and Pew-Thian Yap were supported in part by the National Institute of Mental Health (R01-MH125479). The team at Boston Children’s Hospital was supported in part by the National Institutes of Health (NIH) grants R01-NS106030, R01-EB031849, and R01-EB019483. Team from UW-Madison would like to acknowledge the NIH grants U54HD090256, R01NS092870, R01EB022883, R01AI117924, R01AG027161, RF1AG059312, P50AG033514, R01NS105646, UF1AG051216, R01NS111022, R01NS117568, P01AI132132, R01AI138647, R34DA050258, and R01AG037639. Erick J. Canales-Rodríguez was supported by the Swiss National Science Foundation, Ambizione grant PZ00P2_185814. Matteo Mancini was funded by the Wellcome Trust through a Sir Henry Wellcome Postdoctoral Fellowship [213722/Z/18/Z].

