## Supplementary Information for "Insights from the IronTract challenge: optimal methods for mapping brain pathways from multi-shell diffusion MRI"

#### Supplementary note 1: Description of the methods used by each team in round 1

For each of the teams that participated in round 1 we list the data they used, the pre-processing steps, the orientation reconstruction method, the tractography method, and the post-processing steps.

***Team 1***

**Data**: HCP; DSI.

**Pre-processing**: Data were denoised using Mrtrix3[1,2]. Gibbs ringing artifacts were removed using Mrtrix3[1,3]. Motion and eddy current corrections were performed in FSL[4,5].

**Orientation Reconstruction**: Fiber orientation distribution functions (ODFs) were estimated using the Robust and Unbiased Model-BAsed Spherical Deconvolution (Rumba-SD) method, with 3D total variation regularization (600 iterations)[6].

**Tractography**: Local probabilistic tractography in DIPY (v.1.0.0)[7,8] (bending angle: 20 degrees, step size: 0.2 mm, probability mass function threshold: 0.1, minimum streamline length: 2 mm). A binary dilation was applied to the injection mask, which was used as a seeding mask, with 5,000 seeds randomly placed per voxel. The tracking mask was defined from the Generalized Fractional Anisotropy map estimated using the RUMBA-SD local reconstruction (GFA >= 0.4).

**Post-processing**: Prior to computing volumes, 10% of streamlines with the lowest Cluster Confidence Index (CCI) were removed[7,9]. To reduce the coverage and remove false positives, an iterative thresholding was applied: 200 steps on the log of the streamline count (200 volumes). Additionally, to increase the coverage a Gaussian filter (sigma=1) was applied to the data. Prior to thresholding, the streamline count map was multiplied by the GFA to amplify white matter voxels.

***Team 2***

**Data**: HCP (using only b=6,000 s/mm$^2$ shell)

**Pre-processing:** No pre-processing steps performed.

**Orientation Reconstruction**: Fiber ODFs were estimated using constrained spherical deconvolution (CSD) (lmax: 6)[10] in DIPY. The fiber response function was estimated by a fractional anisotropy (FA) threshold of 0.7 and in a ROI with a radius of 30 centered at the volume center.

**Tractography:** Local probabilistic streamlines tractography in DIPY[7,8] (maximum bending angle: 30 degrees, step size: 0.5 mm, maximum length: 130 mm). The injection mask was binarized and 50,000 seeds were randomly placed.

**Post-processing:** Inclusion regions of interest (ROIs) from the PennCHOP macaque atlas[11] were used to filter the tractography results. These ROIs were selected on the basis of general knowledge of projections of the prefrontal cortex[12] and included: the cingulum bundle, the genu of the corpus callosum, the external capsule, the anterior limb of the internal capsule, and the uncinate fasciculus. Only streamlines intersecting at least one of these ROIs were retained.

***Team 3***

**Data**: HCP.

**Pre-processing**: The data were pre-processed using steps proposed in DESIGNER[13]. Data were denoised[2], corrected for Gibbs ringing artifacts[3] and B1 field inhomogeneity[14] in MRtrix3. Rician bias correction was also performed[15].

**Orientation Reconstruction**: Multi-shell multi-tissue CSD (MSMT-CSD) was performed in MRtrix3[16,17].

**Tractography**: Local probabilistic tractography was performed in MRtrix3[18] (step size: 0.35; minimum length: 20 mm; maximum length: 50,000 mm; trials: 10,000). The seed region was binarized without any additional filtering. The angle threshold was varied in the range 10:10:90 degrees, generating 9 different tractograms. Additional masks were manually drawn to exclude looping streamlines.

**Post-processing**: For each tractogram, different tract density images (TDIs) were created by varying the threshold in the range 0.1:6. For each of the threshold, the TDIs obtained from tractograms generated at different angle thresholds were averaged (irontractchallenge/ITCP2019.sh at main · nadluru/irontractchallenge (github.com)). The averaged TDIs were uploaded to the QMENTA platform for scoring.

***Team 4***

**Data:** HCP; DSI (only first 305 volumes)

**Pre-processing:** Data were denoised with MRtrix3 (v0.3.16)[2] and corrected for motion with ANTs (v2.2.0).

**Orientation Reconstruction:** Fiber ODFs were generated using an in-house implementation of the algorithm in[19] that is based on fitting a three-compartment model on the given data. For both HCP and DSI, eighth order spherical harmonics representation was used.

**Tractography:** Parallel transport tractography (PTT) algorithm[19], implemented in Trekker (https://dmritrekker.github.io/) was used (minRadiusOfCurvature: 0.2 mm; stepSize: 0.01 mm; minLength: 10 mm; maxLength: 100 mm; probeRadius: 0 mm; probeCount: 4; probeQuality: 4; seed_count: 500000). Provided injection site was binarized (>0), manually expanded into the white-matter and used as seed region**.** The probeLength was varied in the range 0.01:0.01:0.15.

**Post-processing:** A 2D ROI was manually drawn on coronal slice 69 to discard connections that pass through these regions. No additional postprocessing was performed.

***Team 5***

**Data**: HCP

**Pre-processing**: No pre-processing steps performed.

**Orientation Reconstruction**: Fiber ODFs were generated using constrained spherical deconvolution (CSD) (Tournier et al., 2007) method in DIPY[7]. The fiber response function was estimated by a fractional anisotropy (FA) threshold of 0.6 and in a ROI with a radius of 30 centered at the volume center.

**Tractography**: Global deterministic shortest-paths tractography was computed using the Cellular Automata Tractography method[20] (https://github.com/andachamamci/CATractography). The injection mask was used as seed region without any modification. The only difference between the method presented in the paper and the method used in the IronTract challenge, is the determination of the anatomical prior (PMAT) for each voxel. Due to the lack of atlas for the macaque brain, an FA mask was used as the anatomical prior, instead of the tissue probability map. The tensor fitting, eigen-decomposition and FA calculation were performed in DIPY package[7]. Then the anatomical priors were set by assigning 1 if the FA was greater than the threshold value of 0.22, and 0 if it was smaller. The FA threshold value was determined by trial on the training dataset.

**Post-processing**: Cellular Automata Tractography produced a map with a connectivity value between each voxel on the image domain and the seed region. The connectivity map was converted to binary maps for submission by using different threshold values. The threshold values were sampled in the range 0.25:0.01:0.45 and 0.45:0.001:0.5, which resulted in 70 total volumes.

***Team 6***

**Data**: HCP.

**Pre-processing**: No pre-processing steps were performed.

**Orientation reconstruction**: Asymmetry spectrum imaging (ASI)[21] was used to allow sub-voxel orientation asymmetry and to take into account a spectrum of hindered/restricted diffusion. To capture white matter configurations such as bending and fanning to mitigate gyral bias, we incorporated information from neighboring voxels to estimate asymmetric fiber ODFs (AFODFs)[22]. We estimated the AFODF at each voxel by enforcing orientation continuity across voxels. Our approach is formulated as a convex problem and does not require initialization with precomputed symmetric FODFs. For visual comparison across approaches in Figure 10, asymmetric FODFs were approximated into symmetric FODFs, but the tractogram was still generated by asymmetric FODFs.

**Tractography**: Local deterministic tractography with asymmetry adaptive forward tracking was performed using the whole brain mask as seeding ROI. The initial step size and initial angle threshold are 0.2 and 60 degrees, respectively. The real step size and angle threshold during the tracing were adaptively updated on the subspace defined by the streamline position, to estimate the forward direction[21,23,24]. No minimum/maximum length were predefined. We seed a fixed number of streamlines 128 per voxel in a mask image. A multi-tissue generalized fractional anisotropy was derived by ASI[21,24] to be used for streamline termination with a fixed threshold of 0.1.

**Post-processing**: We performed a radial search (radius = 4 mm) from each streamline endpoint to locate whether the streamline was close to the injection region. Streamlines that had endpoints close to the injection region were accepted. Voxels with less than 2 streamlines were discarded before converting the tractography sets to binary masks.

***Team 7***

**Data**: HCP

**Pre-processing**: Data were denoised[2] and corrected for Gibbs ringing artifacts and eddy-current distortions in MRtrix3 (*MRtrix3 3.0_RC3-163-ga7ef8ac2*)[1] and FSL (6.0). The preprocessed data were then processed with AMICO (*1.0.2*) to reconstruct the NODDI parameter maps, using a mask computed with MRtrix3. The pre-processed data were further bias-corrected using ANTs *(1.9).*

**Orientation reconstruction**: Fiber ODFs were estimated using MSMT-CSD[1,16,17]. The orientation dispersion (OD) map was thresholded between 0.1 and 0.7 and then binarized and used as a white matter mask for fiber ODFs reconstruction.

**Tractography:** Local probabilistic tractography was performed in MRtrix3 using a dilated version of the injection mask as seeding ROI. As additional inclusion and exclusion criteria, a grey matter (GM) mask and a cerebral spinal fluid (CSF) mask were used. The GM mask was computed by thresholding the intra-cellular volume fraction (ICVF) map below 0.95 and then binarizing it. The CSF mask was computed by binarizing the isotropic volume fraction (ISO-VF). To compensate for the underestimation of long-range connections, the streamlines were reconstructed in two steps using two different minimum streamline thresholds of 10 mm and 50 mm, selecting 10000 streamlines at each step and then combining the resulting streamlines. A range of 30 angular thresholds (from 15 to 44 degrees) was used to generate the different tractography results.

**Post-processing:** Voxels with less than 2 streamlines were discarded before converting the tractography sets to binary masks using MRtrix3.

The reconstruction and tracking procedures used by Team 7 can be reproduced using the TRAMPOLINO *(0.1.3dev)* python package running this single command:

```
trampolino -r results -n mrtrix_workflow recon -i dwi.nii.gz -v grad.txt -b
bvals.txt --opt bthres:0,mask:wm_mask.nii.gz mrtrix_msmt_csd track -s
inject_dil.nii.gz --opt nos:10000,include:gm_mask.nii.gz,csf_mask.nii.gz --angle
15,45 --angle_range --min_length 10,50 --ensemble min_length mrtrix_tckgen
```

***Team 8***

**Data**: HCP

**Pre-processing**: No pre-processing steps performed.

**Orientation Reconstruction:** The diffusion data were reconstructed using generalized q-sampling imaging[25] in DSI Studio (2019 version) with a diffusion sampling length ratio of 0.4.

**Tractography:** A local deterministic fiber tracking algorithm was used[26] (angular threshold: 65 degrees; step size: randomly selected from 0.5 voxel to 1.5 voxels (DSI Studio default setting); minimum length: 10; maximum length: 300 mm). Tractography was seeded inside a whole-brain mask (DSI Studio default setting). The anisotropy threshold was randomly selected (DSI Studio default setting). A total of 50000 tracts were calculated. The injection region is enlarged by 3 times (a volume size of 2.4e+02 mm cubic.) and used as an inclusion ROI. A white-matter mask was created. A termination region was placed at cerebral_white_matter (47,55,47) with a volume size of 3.2e+05 mm cubic. This limits tractography to white matter. The white-matter mask was created using DSI Studio's nonlinear registration**.**

**Post-processing**: Two iterations of topology-informed pruning were used to eliminate false tracts[27]. Tracts were converted to density maps. 40 different thresholds from 0.2 maximum to 0 were used to generate 40 masks for the final results.

Parameter_id to reproduce the results in DSI Studio is c9A99193F6C61D83Edb2041b964350C3cb01dcba

***Team 9***

**Data**: HCP

**Pre-processing**: Data were denoised[2] and corrected for Gibbs ringing artifacts in MRtrix3 (3.0_RC3) [1,3]. Intensity correction was performed in MRtrix and motion/eddy-current correction in FSL (v6.0).

**Orientation Reconstruction**: ODFs were reconstructed using radial DSI (RDSI)[28] in Matlab (R2017a) (https://bitbucket.org/sbaete/rdsi_recon) and fiber directions identified using ODF-Fingerprinting[29] in Matlab (R2017a). Code available at https://bitbucket.org/sbaete/odffingerprinting.

**Tractography**: Local deterministic tractography was performed in DSI Studio (Nov 21, 2018) (Random seeding, step size: 1.5 mm; minimum length: 10 mm; maximum length: 50 mm; 100000 tracts generated from maximum 10 million seed points; streamlines smoothed with 30% of the previous direction). The injection mask was enlarged (4x dilate with MRtrix3) and used as a seed. The turning-angle and FA threshold were varied to produce the different points along the ROC (respectively in the range 40:2:70 and 0.11– 0.7).

**Post-processing:** No post-processing steps were applied.

**Data**: RDSI; 6 datapoints on each of 59 radially outward lines. b-values chosen at equidistant q (b=1025,4099,9223,16396,25618,36890).

**Pre-processing**: Data were denoised[2] in MRtrix3 and corrected for Gibbs ringing artifacts in MRtrix3[1,3]. Intensity correction was performed in MRtrix and motion/eddy-current correction in FSL (v6.0).

**Orientation Reconstruction**: ODFs were reconstructed using RDSI[28] in Matlab (R2017a) (https://bitbucket.org/sbaete/rdsi_recon) and fiber directions identified using ODF-Fingerprinting[29] in Matlab (R2017a). Code available at https://bitbucket.org/sbaete/odffingerprinting.

**Tractography**: Local deterministic tractography was performed in DSI Studio (Nov 21, 2018) (Random seeding, step size: 1.5 mm; fiber lengths > 10mm and < 50mm, 100000 tracts generated from maximum 10 million seedpoints; streamlines smoothed with 30% of the previous direction). The injection mask was enlarged (4x dilate with MRtrix3) and used as a seed. The turning-angle and FA threshold were varied to produce the different points along the ROC (respectively in the range 40:2:70 and 0.11– 0.7).

**Post-processing:** No post-processing steps were applied.

***Team 10***

**Data**: HCP

**Pre-processing**: Data were denoised[2] in MRtrix3 and corrected for N4 bias field in ANTS[14].

**Orientation Reconstruction**: fiber ODFs were reconstructed using MSMT-CSD[16,17].

**Tractography**: Local probabilistic tractography in MRtrix3. Injection site was used as seeding mask and 1 million streamlines were selected. **Tractography**: Local probabilistic tractography in MRtrix3[18]. Injection site was used as seeding mask and 1M streamlines were selected.

**Post-processing**: 3 parameters were varied: the step size ([0.35; 0.7]), the maximum angle threshold ([30°; 45°; 60°]), and the fODF cutoff ([0.05; 0.1; 0.15]).

***Team 11***

**Data**: HCP

**Pre-processing**: Data were corrected for motion and eddy-current distortions[5].

**Orientation reconstruction:** The diffusion data were reconstructed using generalized q-sampling imaging (GQI)[25] in DSI Studio (v1.0) with a diffusion sampling ratio = 1.25. An 8-fold ODF tessellation was used. For each voxel, a maximum of 5 fiber directions were resolved. The diffusion scheme was resampled to ensure balance in 3D space. The order of the x-y-z coordinates in the b-table was preserved.

**Tractography**: A local deterministic fiber tracking algorithm was used[26]. The interpolation was trilinear, and the tracking algorithm was "Streamline (Euler)". Tractography was seeded inside a whole-brain mask and the site of injection was used as inclusion ROI, without any prior modification (quantitative anisotropy (QA) threshold: 0.05; step size: 0.07 mm; default Otsu's threshold:0.6.; max length:900 mm; min length: 10 mm). The seed orientation was primary, the seed position was sub voxel, the randomize seeding parameter was on. The fiber trajectories were smoothed by averaging the propagation direction with a percentage of the previous direction (randomly selected from 0% to 95%.). A total of 50,000 tracts were generated. The angular threshold was varied from 22 to 90 degrees with a step size of 2 degrees, for a total of 35 tractography reconstructions.

**Post-processing:** Tracks were converted to density maps. To ensure that these volumes had the same orientation as the original DWIs (radiological), they were re-oriented using FSL. In the case of the tractography reconstruction of the validation data, the tracts masks were further registered to the first volume of the DWI image, by using FSL FLIRT, and then thresholded (0.9) and binarized with fslmaths. No additional masks were used to constrain tractography.

***Team 12***

**Data**: HCP

**Pre-processing**: Each of the volumes was denoised independently using the BM4D algorithm[30]. Skull removal: A volume mask was generated by simple thresholding of the b0 image. The boundary of this mask was used to generate a skull mask using Maurer distance transform.

**Orientation Reconstruction**: Fiber ODFs were reconstructed using spherical deconvolution[31].

**Tractography**: Local deterministic tractography was performed using the EuDX algorithm[32] in DIPY 0.16[7]. A step size of 0.50 mm was used. The stopping criterion was generalized fractional anisotropy of less than 0.20. A total of 10 tractograms were obtained by varying the maximum angle threshold between 20 and 40 degrees.

**Post-processing**: Tractograms were converted to density maps and binarized (> 0.1).

**Supplementary note 2: Description of the methods used by each team in round 2**

For each of the teams that participated in round 2 we list the data they used, the orientation reconstruction method, and the tractography method. The pre- and post-processing steps were the same for all teams in round 2, as described in the Methods section, and are therefore omitted here. Returning teams that used the same method they used in round 1 are not listed here.

***Team 12***

**Data**: HCP

**Orientation Reconstruction**: Fiber ODFs were reconstructed using a deep learning-based approach[33]. To estimate the fiber ODF in each voxel, the model used the diffusion data in the 3*3*3-voxel neighborhood around that voxel. The measurements were represented in order-4 spherical harmonic coefficients. These coefficients were used as the input to the model. The model output was the ODF, represented in order-8 spherical harmonics. Training data were generated using the IronTract training scan. Voxels with generalized fractional anisotropy of 0.20 and above were considered for training. For these voxels, Constrained Spherical Deconvolution was applied to generate the reference ODFs. The network first projected the spherical harmonic representation of each of the 27 voxels in the neighborhood onto a lower dimension of 10. These representations were concatenated to form a vector of length 270. This vector went through a succession of 5 fully connected hidden layers with ReLU activations. The final (output) layer did not have a ReLU activation to allow for negative spherical harmonic coefficients. The model was trained by minimizing the L2 norm between the target ODF and predicted ODF. The trained model was then applied in the same fashion on the validation scan to estimate the ODF for each voxel.

**Tractography**: Local deterministic tractography was performed using the EuDX algorithm[32] in DIPY version 0.16[7]. A step size of 0.50 mm was used. The stopping criterion was generalized fractional anisotropy of less than 0.20. Maximum angle between steps was varied between 20 and 40 degrees.

***Team 13***

**Data**: HCP

**Orientation Reconstruction**: fiber ODFs were reconstructed using CSD[10] in DIPY[7]. The CSD fiber response function is estimated using an FA threshold of 0.6 and a ROI radius of 30 degrees at the volume center. The pipeline is available in CATractography code[20] (https://github.com/andachamamci/CATractography).

**Tractography**: First, a graph based on fiber ODFs is constructed, on which the shortest-paths are computed[20]. Next, using the edge weights as rewards, the value for each action (Q-function) is computed by a training process in reinforcement learning context[34]. Finally, probabilistic streamline tractography is run on the Q-function, setting the maximum bending to 45 degrees, as implemented in DIPY. The injection mask was used as a seed mask without any modification. Probabilistic tracking is initialized for 64,000 seeds per voxel in the seed mask. The anatomical prior was obtained by using an FA mask with a threshold of 0.22. The FA threshold value is determined by trial on the training dataset

***Team 14***

**Data**: HCP

**Orientation Reconstruction**: Fiber ODFs were estimated using CSD[10] using implementation from ExploreDTI[35] available via MRIToolkit (https://github.com/delucaal/MRIToolKit).

**Tractography**: A deterministic multi-level fiber tractography algorithm was used[36] with 2-level reconstruction (FOD peak threshold: 0.01, angular threshold: 45°). The mask provided with the post-processing *Anatomical ROIs* script was used as a seed region. Each voxel was subsampled using 2x2x2 uniform greed. The injection point was used as a target. Exclusion ROIs were manually set to remove streamlines traversing to the hemisphere contralateral to the injection point (except for those close to target ROI).

**Data**: DSI

**Orientation Reconstruction**: Fiber ODFs were estimated with the generalized Richardson-Lucy (GRL) spherical deconvolution method[37] using MRIToolkit.

**Tractography**: Local deterministic multi-peak tractography[38] was used (FOD peak threshold: 0.01, angular threshold: 45°, step-size 0.35 mm) (https://github.com/delucaal/MRIToolKit). Seeds were uniformly sampled at 0.35 $mm^3$ isotropic resolution in the injection mask. NOT gates were manually placed to remove streamlines traversing to the hemisphere contralateral to the injection point (except for those close to target ROI).

### Supplementary note 3: Additional analyses

#### Bundle-wise TPR.

As an alternative to the voxel-wise TPR (*see Methods, ROC analysis*), we also investigated how many of the main white-matter areas that were included in the tracer mask were reached by each tractography method. The goal was to determine what FPR we would have to tolerate with each method to reach the main bundles that the injection site projects to, and what tractography threshold would allow us to achieve that. To this end, every voxel included in the tracer mask in dMRI space was labeled by AY and CM. For the training case, voxels were assigned to one of 8 classes: anterior frontal white matter (AF); anterior limb of the internal capsule (ALIC); cingulum bundle (CB); corpus callosum (CC); external capsule (EC); medial prefrontal white matter (MPF); lateral prefrontal white matter (LPF); uncinate fasciculus (UF). For the validation case, voxels were assigned to one of 10 classes: ALIC; brainstem fibers (BS); commissural fibers (CF); CB; CC; EC; extreme capsule (EmC); LPF; thalamic fibers (ThF); UF. We assumed that tractography reached one of the above labels successfully if it reached at least 50% of the voxels in the label. We computed the *bundle-wise* TPR, which we defined as the percentage of labels reached successfully by each tractogram. We then identified the tractogram threshold at which each submission achieved a bundle-wise TPR of 0.8, *i.e.,* reached 80% of the white-matter regions that the injection site projects to. The goal was to examine if there was a similar threshold for which most methods achieved satisfactory coverage of the true bundles. If such a common threshold exists, it may be a sensible choice for users of tractography, in the general scenario where ground truth is not available.

Results of these additional analyses are shown in Supplementary figures S4 and S5. Figure S4 shows ROC curves for round 2 with the bundle-wise TPR, *i.e.,* the portion of white-matter regions where each submission achieved at least 50% coverage. For this and subsequent results presented in this section, we show only one submission per team (the one that achieved the highest AUC score on the validation case). Results are shown for post-processing by a Gaussian filter. Some of the submissions that used deterministic tractography (Team 8 for HCP and DSI schemes; Team 12 for HCP scheme) could not reach all the ten white-matter regions and are thus not shown. There was considerable variability across submissions, with Teams 1 and 2 reaching 50% coverage of all the regions with FPR < 0.11, and the remaining submissions with FPR > 0.15. Deterministic methods (solid lines) operate at lower specificity levels, and are only able to reach all regions at the cost of FPR > 0.22. In some cases, submissions that used the same orientation reconstruction method (M-CSD[16,17]) achieved coverage of all regions at very different FPR levels, suggesting that other algorithmic choices had an impact on the TPR/FPR trade-off. It is worth noting that for the DSI scheme, all submissions (including deterministic) reached all ten regions at higher specificity levels (FPR $\leq$ 0.15) than for the HCP scheme, and that the submission that used Rumba-SD was able to reach 9 out of 10 regions with an FPR as low as 0.06.

Figure S5 shows the tractogram threshold for which each team achieved bundle-wise TPR = 0.8. Overall, most submissions needed very relaxed thresholds (< 0.02 of the maximum value in the tractogram). Only one submission, using ASI[21] and deterministic tractography[24], achieved this coverage at a much more stringent threshold (0.13 of the maximum value in the tractogram). However, this submission also produced a much higher FPR at that threshold. For most submissions, a slightly higher (more stringent) threshold was needed in the validation case than the training case.

**Hausdorff distance.**

As an alternative error metric to the FPR (*see Methods, ROC analysis*), we computed the modified Hausdorff distance (MHD)[39] between the tracer mask and the tractogram. The MHD between two set of points S and T is defined as the minimum distance between a point in one set and any point in the other set, averaged over all points in the two sets:

$$MHD(S,T) = \frac{1}{|S|}\sum_{s\epsilon S} \quad min_{t\epsilon T} d(s,t) + \frac{1}{|T|}\sum_{t\epsilon T} \quad min_{s\epsilon S} d(t,s),$$

where d(·,·) is the Euclidean distance between two points, and |·| is the size of a set. Greater MHD indicates greater deviation of the tractography volume from the tracer.

Results of these additional analyses are shown in supplementary figures S6. The MHD between the tracer mask and the tractogram is plotted against the TPR. While the FPR penalizes all FPs equally, the MHD measures how far from the tracer mask the FPs occur. The plots show that the MHD was greater for the validation than the training case for all submissions, even those that achieved similarly high AUC score in the two cases. At the same sensitivity level, MHD was greater for deterministic than probabilistic methods. Similarly to what we observed with the bundle-wise ROCs of Figure S4, there were submissions that used the same orientation reconstruction method (CSD) but had very different MHDs at the same level of sensitivity (3-8 mm range at TPR = 0.8). The MHD was below 10 mm for all submissions and all levels of sensitivity.

**Analysis of orientation peaks**

For each submission, we extracted the orientation dispersion and the peak orientation for a maximum of 3 peaks per orientation distribution function (ODF) in MRtrix3[40,41]. Only peaks larger than a threshold (0.2 times the largest ODF value) were selected, as very small peaks would typically not be used in tractography. We computed the angular difference between the first and second peak of each submission and the peaks of the submission that achieved the highest performance across the two rounds (Team 1, Rumba-SD DSI). Angular differences were computed separately for the first and second largest peak, and averaged over voxels in each of the ROIs.

Results of this analysis are presented in supplementary figure 7. Results showed that, although the peak orientation was not the only factor that determined accuracy, submissions that had smaller differences in AUC with Team 1 (< 20%) also had smaller angular differences (< 15°). However, there were also teams with similar peak orientations that achieved much lower AUC scores.

**Supplementary Tables**

| Submission Number | Team Number | Acquisition Scheme | Orientation Reconstruction Method | Propagation Method | Tractography Algo. | AUC | TPR |
|---|---|---|---|---|---|---|---|
| 01 | 1 | HCP | Rumba-SD | Probabilistic | DIPY_prob | 0.2468 | 0.816484 |
| 02 | 2 | HCP | CSD | Probabilistic | DIPY_prob | 0.2365 | 0.792842 |
| 06 | 6 | HCP | ASI | Deterministic | iFOD2 | 0.1819 | 0.540728 |
| 03 | 3 | HCP | M-CSD | Probabilistic | Trekker_PPT | 0.1812 | 0.551639 |
| 04 | 4 | HCP | 3CMP | Probabilistic | CAT | 0.1732 | 0.619192 |
| 07 | 7 | HCP | M-CSD | Probabilistic | AAFT | 0.1563 | 0.422622 |
| 05 | 5 | HCP | CSD | Deterministic | iFOD2 | 0.1504 | 0.443055 |
| 09 | 9 | HCP | RDSI | Deterministic | DSI_Studio | 0.1401 | 0.49053 |
| 08 | 8 | HCP | GQI | Deterministic | DSI_Studio | 0.1252 | 0.481399 |
| 10 | 10 | HCP | M-CSD | Probabilistic | iFOD2 | 0.0945 | 0.210048 |
| 11 | 11 | HCP | GQI | Deterministic | DSI_Studio | 0.0931 | 0.186954 |
| 12 | 12 | HCP | RL | Deterministic | DIPY_EuDX | 0.0058 | 0.773927 |
| 13 | 1 | DSI | Rumba-SD | Probabilistic | DIPY_prob | 0.2667 | 0.96502 |
| 16 | 9 | DSI | RDSI | Deterministic | Trekker_PPT | 0.1846 | 0.56446 |
| 14 | 4 | DSI | 3CMP | Probabilistic | DSI_Studio | 0.1669 | 0.475926 |
| 15 | 8 | DSI | GQI | Deterministic | DSI_Studio | 0.1294 | 0.474922 |

**Supplementary Table 1. Round 1 submission details.** For each submission, the table reports the team number, acquisition scheme, orientation reconstruction method, propagation method (probabilistic or deterministic tractography), tractography algorithm, the AUC score, and the TPR at FPR = 0.1 that were achieved on the validation case. Orientation reconstruction methods: ASI = asymmetry spectrum imaging[22]; 3CMP = three compartment model[42]; CSD = constrained spherical deconvolution[10]; ; GQI = generalized Q-ball imaging[25]; M-CSD = multi-shell multi-tissue CSD; RDSI = radial diffusion spectrum imaging[28,29]; RL = Richardson Lucy[31]; Rumba-SD = robust and unbiased model-based spherical deconvolution[6]. Tractography algorithms: AAFT = asymmetry adaptive forward tracking[21,24]; CAT = cellular automata tractography[20]; DIPY_Prob = DIPY probabilistic[7,8]; DIPY_EuDX = DIPY EuDX[32]; DSI_Studio[26]; iFOD2 = Second-order Integration over Fiber Orientation Distributions[18]; Trekker_PPT = Trekker parallel transport tractography[19].

| Submission number | Team | Acquisition Scheme | Orientation Reconstruction method | Post-processing | Propagation Method | Tractography Algo. | AUC | TPR |
|---|---|---|---|---|---|---|---|---|
| 17 | 01 | HCP | Rumba-SD | Gaussian Filter | Probabilistic | DIPY_prob | 0.2456 | 0.826684 |
| 18 | 01 | HCP | Rumba-SD | Anatomical ROIs | Probabilistic | DIPY_prob | 0.2554 | 0.901975 |
| 19 | 01 | DSI | Rumba-SD | Gaussian Filter | Probabilistic | DIPY_prob | 0.2659 | 0.960799 |
| 20 | 01 | DSI | Rumba-SD | Anatomical ROIs | Probabilistic | DIPY_prob | 0.2669 | 0.956969 |
| 21 | 01 | HCP | CSD | Gaussian Filter | Probabilistic | DIPY_prob | 0.2455 | 0.861219 |
| 22 | 01 | HCP | CSD | Anatomical ROIs | Probabilistic | DIPY_prob | 0.2503 | 0.880231 |
| 23 | 01 | DSI | DSI | Gaussian Filter | Probabilistic | DIPY_prob | 0.2275 | 0.7819 |
| 24 | 01 | DSI | DSI | Anatomical ROIs | Probabilistic | DIPY_prob | 0.2427 | 0.837326 |
| 25 | 02 | HCP | CSD | Gaussian Filter | Probabilistic | DIPY_prob | 0.2517 | 0.901674 |
| 26 | 02 | HCP | CSD | Anatomical ROIs | Probabilistic | DIPY_prob | 0.257 | 0.924062 |
| 27 | 02 | HCP | CSD | Gaussian Filter | Probabilistic | DIPY_prob | 0.2538 | 0.920565 |
| 28 | 02 | HCP | CSD | Anatomical ROIs | Probabilistic | DIPY_prob | 0.2583 | 0.919389 |
| 29 | 02 | DSI | DSI | Gaussian Filter | Probabilistic | DIPY_prob | 0.2538 | 0.920565 |
| 30 | 02 | DSI | DSI | Anatomical ROIs | Probabilistic | DIPY_prob | 0.2583 | 0.919389 |
| 31 | 03 | HCP | M-CSD | Gaussian Filter | Probabilistic | MRtrix3_iFOD 2 | 0.2008 | 0.58796 |
| 32 | 03 | HCP | M-CSD | Anatomical ROIs | Probabilistic | MRtrix3_iFOD 2 | 0.2286 | 0.717096 |
| 33 | 04 | HCP | 3CMP | Gaussian Filter | Probabilistic | Trekker_PPT | 0.2016 | 0.58863 |
| 34 | 04 | HCP | 3CMP | Anatomical ROIs | Probabilistic | Trekker_PPT | 0.2232 | 0.715329 |
| 35 | 04 | DSI | 3CMP | Gaussian Filter | Probabilistic | Trekker_PPT | 0.2249 | 0.716322 |
| 36 | 04 | DSI | 3CMP | Anatomical ROIs | Probabilistic | Trekker_PPT | 0.2528 | 0.909865 |
| 37 | 04 | HCP | M-CSD | Gaussian Filter | Probabilistic | Trekker_PPT | 0.199 | 0.603108 |
| 38 | 04 | HCP | M-CSD | Anatomical ROIs | Probabilistic | Trekker_PPT | 0.2163 | 0.675006 |
| 39 | 04 | HCP | 3CMP | Gaussian Filter | Probabilistic | Trekker_PPT | 0.2153 | 0.672953 |

| | | | | | | | | |
|---|---|---|---|---|---|---|---|---|
| 40 | 04 | HCP | 3CMP | Anatomical ROIs | Probabilistic | Trekker_PPT | 0.2316 | 0.772273 |
| 41 | 04 | DSI | 3CMP | Gaussian Filter | Probabilistic | Trekker_PPT | 0.1855 | 0.534942 |
| 42 | 04 | DSI | 3CMP | Anatomical ROIs | Probabilistic | Trekker_PPT | 0.2112 | 0.618411 |
| 43 | 05 | HCP | CSD | Gaussian Filter | Deterministic | CAT | 0.1952 | 0.646196 |
| 44 | 06 | HCP | ASI | Gaussian Filter | Deterministic | AAFT | 0.1602 | 0.419013 |
| 45 | 06 | HCP | ASI | Anatomical ROIs | Deterministic | AAFT | 0.2196 | 0.729133 |
| 46 | 07 | HCP | M-CSD | Gaussian Filter | Probabilistic | iFOD2 | 0.2066 | 0.619101 |
| 47 | 07 | HCP | M-CSD | Anatomical ROIs | Probabilistic | iFOD2 | 0.2167 | 0.682211 |
| 48 | 08 | HCP | GQI | Gaussian Filter | Deterministic | DSI_Studio | 0.1184 | 0.374907 |
| 49 | 08 | HCP | GQI | Anatomical ROIs | Deterministic | DSI_Studio | 0.1225 | 0.390389 |
| 50 | 08 | DSI | GQI | Gaussian Filter | Deterministic | DSI_Studio | 0.116 | 0.358561 |
| 51 | 08 | DSI | GQI | Anatomical ROIs | Deterministic | DSI_Studio | 0.1192 | 0.366447 |
| 52 | 09 | HCP | RDSI | Gaussian Filter | Deterministic | DSI_Studio | 0.1682 | 0.482016 |
| 53 | 09 | HCP | RDSI | Anatomical ROIs | Deterministic | DSI_Studio | 0.1876 | 0.572518 |
| 54 | 09 | DSI | RDSI | Gaussian Filter | Deterministic | DSI_Studio | 0.2156 | 0.743257 |
| 55 | 09 | DSI | RDSI | Anatomical ROIs | Deterministic | DSI_Studio | 0.2214 | 0.778449 |
| 56 | 10 | HCP | M-CSD | Gaussian Filter | Probabilistic | iFOD2 | 0.1911 | 0.54878 |
| 57 | 10 | HCP | M-CSD | Anatomical ROIs | Probabilistic | iFOD2 | 0.2113 | 0.596767 |
| 58 | 11 | HCP | GQI | Gaussian Filter | Deterministic | DSI_Studio | 0.1704 | 0.466862 |
| 59 | 11 | HCP | GQI | Anatomical ROIs | Deterministic | DSI_Studio | 0.1746 | 0.485227 |
| 60 | 12 | HCP | ML | Gaussian Filter | Deterministic | DIPY_EuDX | 0.1518 | 0.497436 |
| 61 | 14 | HCP | CSD | Anatomical ROIs | Deterministic Probabilistic | MLFT | 0.1643 | 0.503821 |
| 62 | 14 | HCP | GRL | Anatomical ROIs | Deterministic | MLFT | 0.1486 | 0.424491 |
| 63 | 14 | HCP | GRL | Anatomical ROIs | Deterministic | MLFT | 0.0954 | 0.301591 |
| 64 | 14 | DSI | GRL | Anatomical ROIs | Deterministic | MLFT | 0.118 | 0.388603 |
| 65 | 13 | HCP | CSD | Gaussian Filter | Probabilistic | Q-routing | 0.2091 | 0.690256 |
| 66 | 13 | HCP | CSD | Anatomical ROIs | Probabilistic | Q-routing | 0.2221 | 0.752335 |

**Supplementary Table 2. Round 2 submission details**. For each round 2 submission, the table reports the team number, acquisition scheme, orientation reconstruction method, propagation method (probabilistic or deterministic tractography), the AUC score, and the TPR at FPR = 0.1 that were achieved on the validation case. Orientation reconstruction methods: ASI = asymmetry spectrum imaging[22]; 3CMP = three compartment model[43]; CSD = constrained spherical deconvolution[10]; DSI = Diffusion spectrum imaging[44]; ; GQI = generalized Q-ball imaging[25]; GRL= generalized Richardson-Lucy[37]; M-CSD = multi-shell multi-tissue CSD[16,17]; ML = machine learning-based reconstruction[33]; RL= Richardson Lucy[31], RDSI = radial diffusion spectrum imaging[28,29]; Rumba-SD = robust and unbiased model-based spherical deconvolution[6]. Tractography algorithms: AAFT = asymmetry adaptive forward tracking[21,24]; CAT = cellular automata tractography[20]; DIPY_Prob = DIPY probabilistic[7,8]; DIPY_EuDX = DIPY EuDX[32]; DSI_Studio[26]; iFOD2 = Second-order Integration over Fiber Orientation Distributions[18]; MLFT = multi-level fiber tractography[36]; Q-routing = Q-routing based method[34]; Trekker_PPT = Trekker parallel transport tractography[19].

### Supplementary Figures

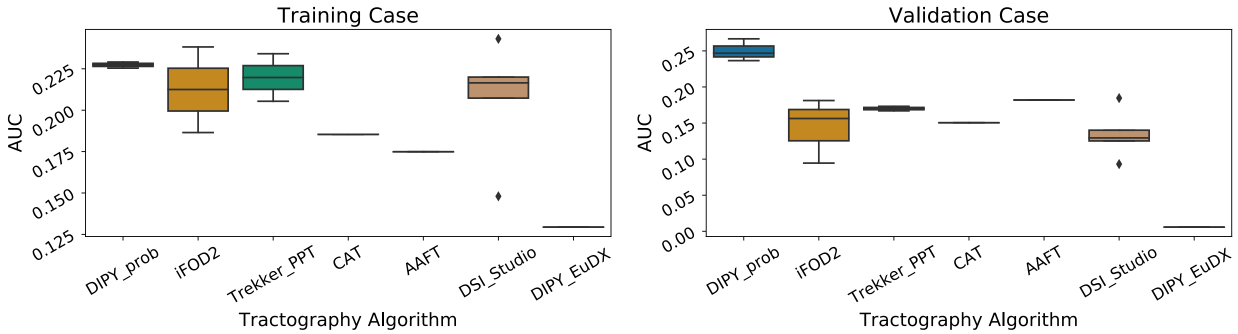


**Supplementary Figure 1**. AUC scores are shown by tractography algorithm for the training case (left) and the validation case (right). AAFT = asymmetry adaptive forward tracking[21,24]; CAT = cellular automata tractography[20]; DIPY_Prob = DIPY probabilistic[7,8]; DIPY_EuDX = DIPY EuDX[32]; DSI_Studio[26]; iFOD2 = Second-order Integration over Fiber Orientation Distributions[18]; MLFT = multi-level fiber tractography[36]; Q-routing =; Trekker_PPT = Trekker parallel transport tractography[19].

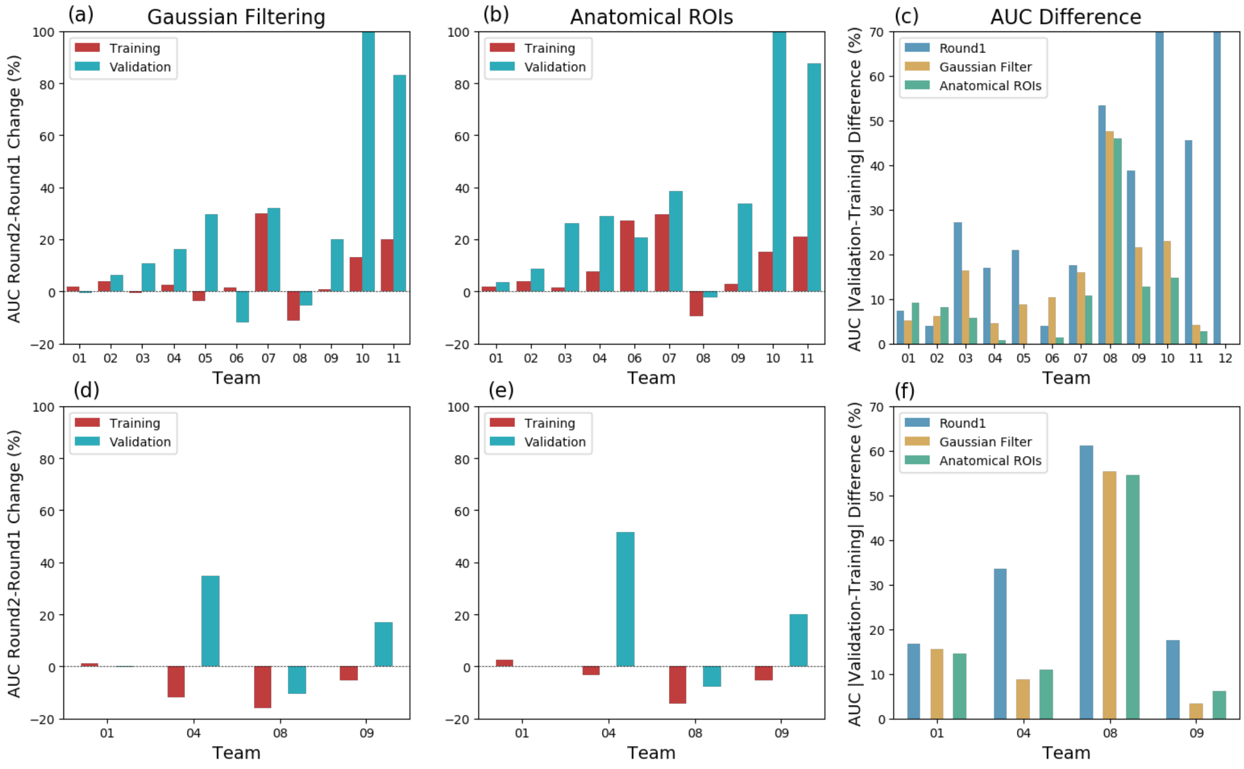


**Supplementary Figure 2**. Top row: HCP acquisition. Bottom row: DSI acquisition. **a,d)** Percent change in AUC scores between round 1 and round 2 submissions for training and validation cases, where round 2 submissions were post-processed with Gaussian Filtering. **b,e)** As above, for round 2 submissions post-processed with anatomical ROIs. **c,f)**

Difference in AUC scores between the training and validation cases, for round 1 and for each of the two post-processing strategies in round 2.

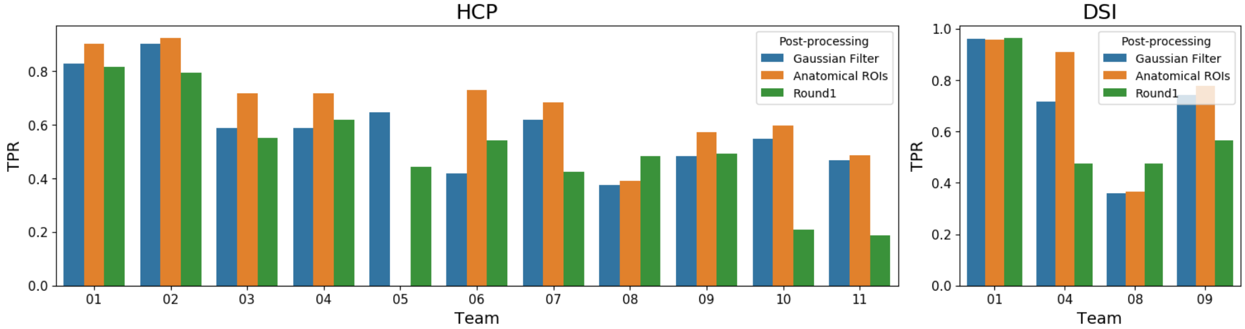


**Supplementary Figure 3.** Bar graph of the TPR achieved for the validation case by each team at FPR=0.1, in round 1 and with each of the two post-processing strategies in round 2 (left: HCP data; right: DSI data).

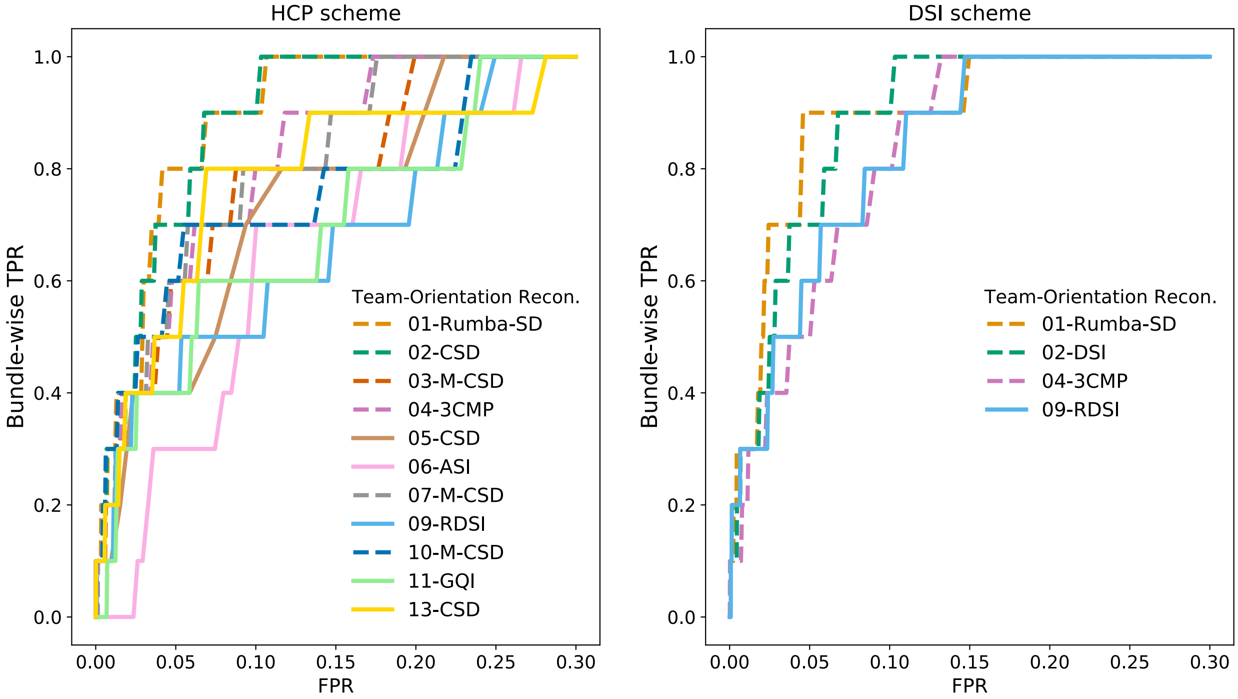


**Supplementary Figure 4. Bundle-wise TPR.** ROC curves with the bundle-wise TPR are shown for the validation case and post-processing by a Gaussian filter. The bundle-wise TPR is defined as the portion of white-matter regions (ALIC, BS, CB, CC, CF, EC, EmC, LPF, ThF, UF) where a submission achieved at least 50% coverage. ASI = asymmetry spectrum imaging[22]; 3CMP = three compartment model[42]; CSD = constrained spherical deconvolution[10]; DSI = Diffusion spectrum imaging[44]; GQI = generalized Q-ball imaging[25]; M-CSD = multi-shell multi-tissue CSD[16,17]; RDSI = radial diffusion spectrum imaging[28,29]; Rumba-SD = robust and unbiased model-based spherical deconvolution[6].

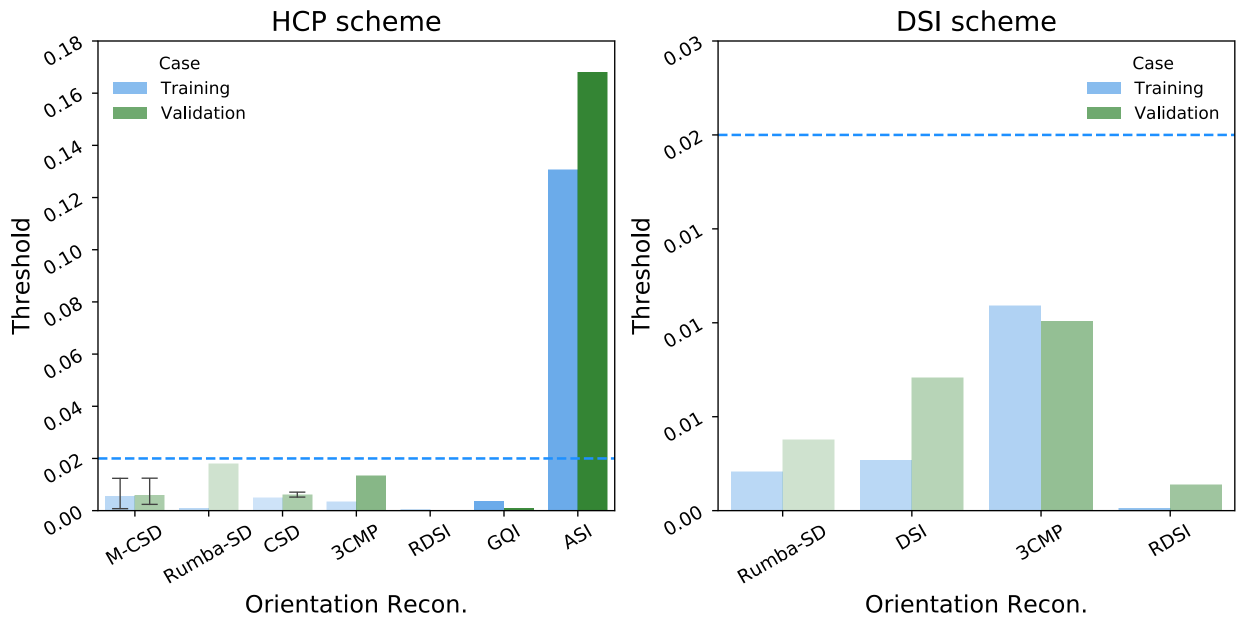


**Supplementary Figure 5. Tractogram thresholds needed to achieve high coverage of the tracer mask.** Bar plots show the tractogram threshold (with respect to the maximum value in the tractogram) for which each submission achieved bundle-wise TPR = 0.8. Submissions are grouped by the orientation reconstruction method that they used. Results are shown for the training (blue) and validation (green) case, and for the HCP (left) and DSI (right) sampling scheme. The bars are ordered along the x-axis by the FPR of the corresponding submissions, which is also indicated by the saturation level of each bar. ASI = asymmetry spectrum imaging[21,22]; 3CMP = three compartment model[42]; CSD = constrained spherical deconvolution[10]; DSI = Diffusion spectrum imaging[44]; GQI = generalized Q-ball imaging[25]; M-CSD = multi-shell multi-tissue CSD[16,17]; RDSI = radial diffusion spectrum imaging[28,29]; Rumba-SD = robust and unbiased model-based spherical deconvolution[6].

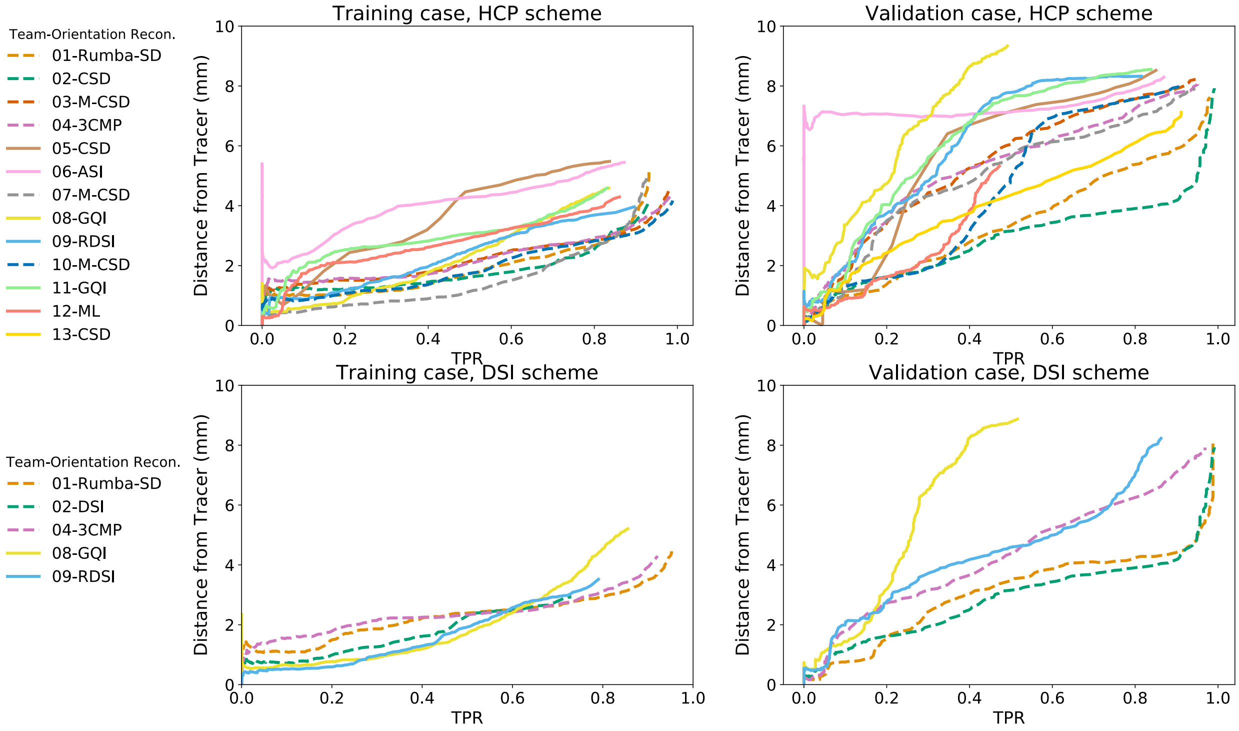


**Supplementary Figure 6. Distance between tractograms and the tracer mask.** The MHD (in mm) between the tractogram and the tracer mask is plotted versus TPR, for the validation case and post-processing by a Gaussian filter. ASI = asymmetry spectrum imaging[21,22]; 3CMP = three compartment model[42]; CSD = constrained spherical deconvolution[10]; DSI = Diffusion spectrum imaging[44]; GQI = generalized Q-ball imaging[25]; M-CSD = multi-shell multi-tissue CSD[16,17]; ML = machine learning-based reconstruction[33]; RDSI = radial diffusion spectrum imaging[28,29]; Rumba-SD = robust and unbiased model-based spherical deconvolution[6].

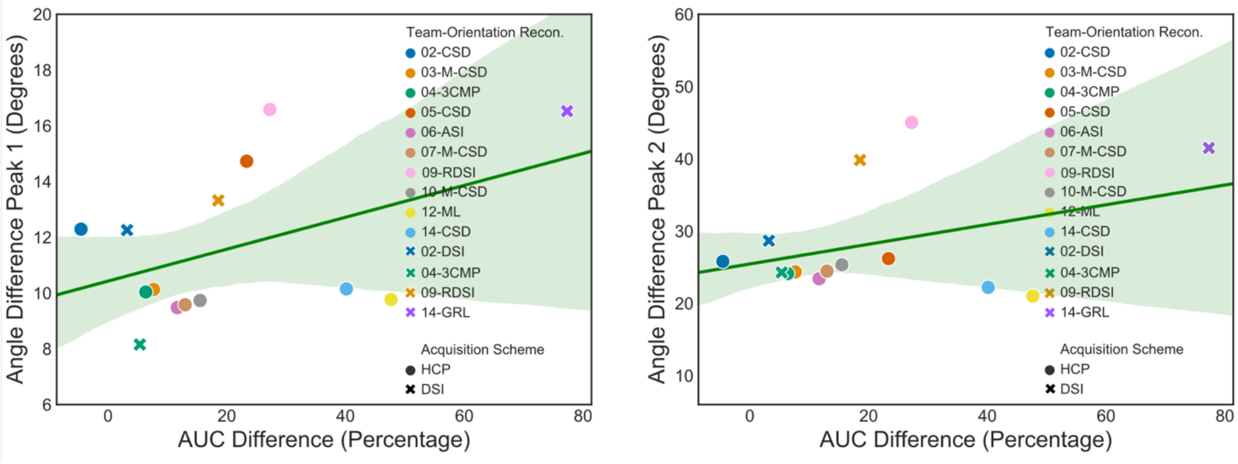


**Supplementary Figure 7**. Scatter plots of the mean peak difference (degrees) and AUC difference (%) between each submission and Team 1 for the validation case. A linear regression model fit is overlaid on the plots, along with a 95% confidence interval band in green. The mean angular difference is shown for the two main peaks (left: peak 1; right: peak 2), averaged over the entire tracer mask. ASI = asymmetry spectrum imaging[21,22]; 3CMP = three compartment model[42]; CSD = constrained spherical deconvolution[10]; DSI = Diffusion spectrum imaging[44]; GQI = generalized Q-ball imaging[25]; M-CSD = multi-shell multi-tissue CSD[16,17]; ML = machine learning-based reconstruction[33]; RDSI = radial diffusion spectrum imaging[28,29]; Rumba-SD = robust and unbiased model-based spherical deconvolution[6].

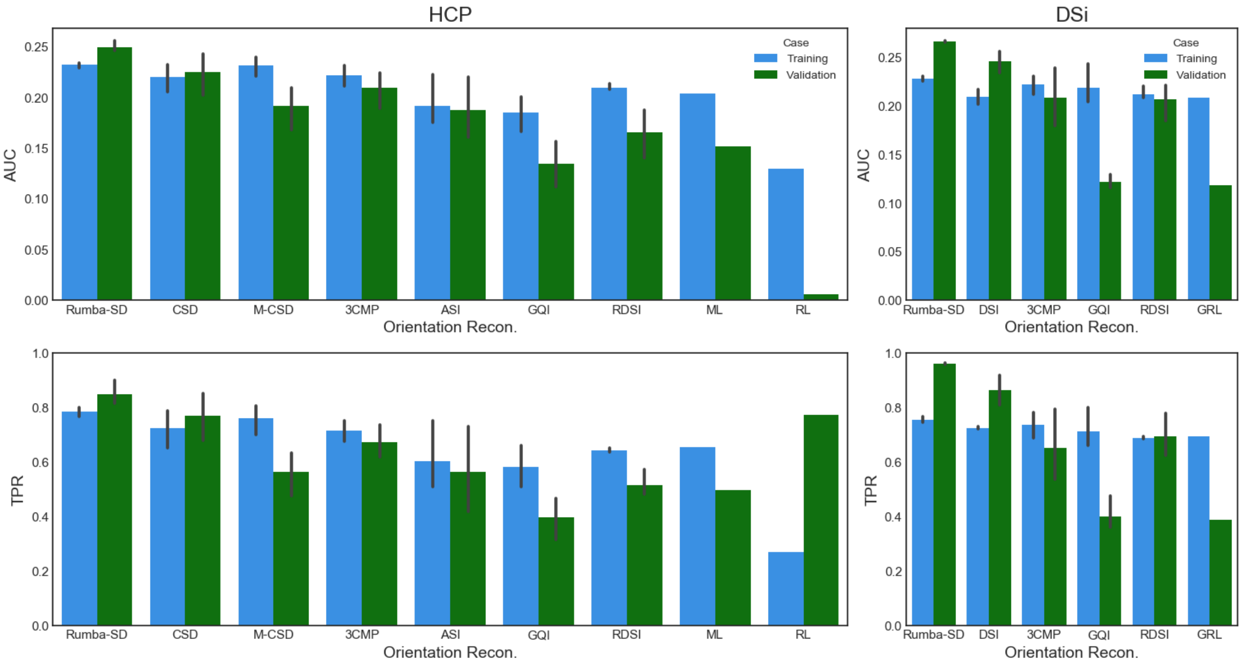


**Supplementary Figure 8. Top)** AUC for validation case for each diffusion model across different submissions for both training and validation case in round 2. **Bottom)** Highest TPR obtained at FPR = 0.1 for the validation case for each orientation reconstruction method across different submissions, for both training and validation case in round 2. ASI = asymmetry spectrum imaging[21,22]; 3CMP = three compartment model[42]; CSD = constrained spherical deconvolution[10]; DSI = Diffusion spectrum imaging[44]; GQI = generalized Q-ball imaging[25]; M-CSD = multi-shell multi-tissue CSD[16,17]; ML = machine learning-based reconstruction[33]; RDSI = radial diffusion spectrum imaging[28,29]; Rumba-SD = robust and unbiased model-based spherical deconvolution[6].
